# Cryo-EM Structures of the SARS-CoV-2 Endoribonuclease Nsp15

**DOI:** 10.1101/2020.08.11.244863

**Authors:** Monica C. Pillon, Meredith N. Frazier, Lucas B. Dillard, Jason G. Williams, Seda Kocaman, Juno M. Krahn, Lalith Perera, Cassandra K. Hayne, Jacob Gordon, Zachary D. Stewart, Mack Sobhany, Leesa J. Deterding, Allen L. Hsu, Venkata P. Dandey, Mario J. Borgnia, Robin E. Stanley

## Abstract

New therapeutics are urgently needed to inhibit SARS-CoV-2, the virus responsible for the on-going Covid-19 pandemic. Nsp15, a uridine-specific endoribonuclease found in all coronaviruses, processes viral RNA to evade detection by RNA-activated host defense systems, making it a promising drug target. Previous work with SARS-CoV-1 established that Nsp15 is active as a hexamer, yet how Nsp15 recognizes and processes viral RNA remains unknown. Here we report a series of cryo-EM reconstructions of SARS-CoV-2 Nsp15. The UTP-bound cryo-EM reconstruction at 3.36 Å resolution provides molecular details into how critical residues within the Nsp15 active site recognize uridine and facilitate catalysis of the phosphodiester bond, whereas the apo-states reveal active site conformational heterogeneity. We further demonstrate the specificity and mechanism of nuclease activity by analyzing Nsp15 products using mass spectrometry. Collectively, these findings advance understanding of how Nsp15 processes viral RNA and provide a structural framework for the development of new therapeutics.

## Introduction

Severe Acute Respiratory Syndrome Coronavirus 2 (SARS-CoV-2) is the virus at the center of the unprecedented Covid-19 global health pandemic. All coronaviruses have large single-stranded, positive-sense RNA genomes that harbor two open reading frames that are translated upon entry into the host (Fig. 1). The two open reading frames in SARS-CoV-2 are translated by host ribosomes into two long polyproteins that are processed by viral proteases into distinct non-structural proteins (designated by the Nsp acronym). These non-structural proteins serve a variety of roles in virus replication, virus assembly, and evasion of host viral sensors, but the precise function of many of these proteins are poorly understood (*1*). Although some inhibitors show promise as antiviral drugs directed specifically against SARS-CoV-2 non-structural proteins, there is a pressing need to understand the structure and function of the non-structural proteins to aid in the development of new and effective antiviral therapies for Covid-19 (*2*).

**Fig.1.**
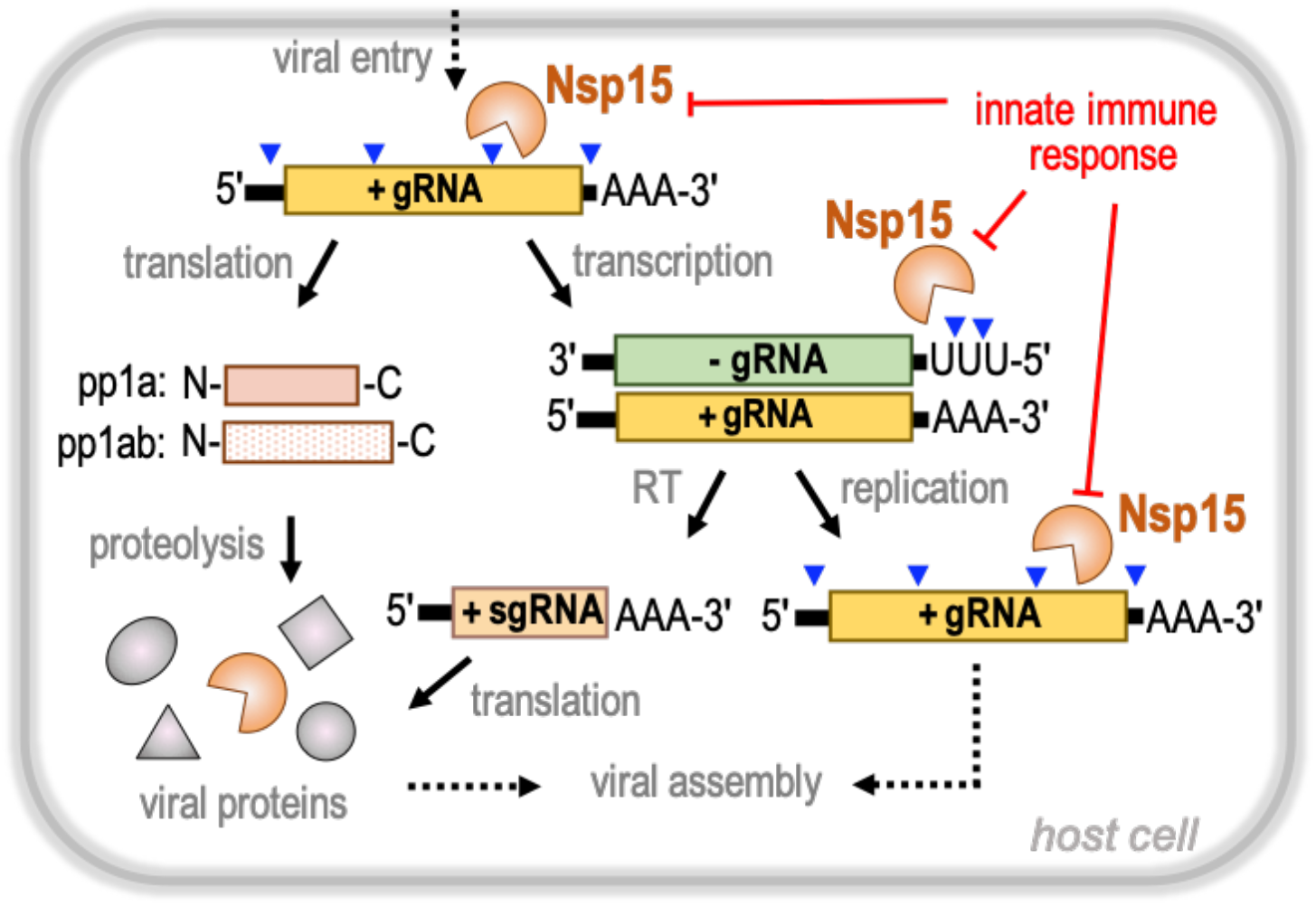
Nsp15 is a central regulator of SARS-CoV-2 RNA processing. Upon viral entry, the single-strand SARS-CoV-2 positive-sense genomic RNA (+gRNA; yellow) is released into the cytoplasm and translated by host ribosomes to generate viral polyproteins pp1a and pp1ab. Subsequent proteolytic cleavage of the polyproteins results in a variety of non-structural proteins (Nsp) essential for diverse viral functions. Transcription of the positive-sense genomic RNA produces a negative-sense genomic RNA (-gRNA; green) intermediate. The negative-sense strand is a template for reverse transcription (RT) generating a series of positive-sense subgenomic RNAs (+sgRNA; brown), which are translated into diverse structural proteins. In addition, the negative-sense strand is also the template for viral replication. The non-structural protein Nsp15 (orange) is a poly-(U) specific endonuclease that cleaves 3′ to uridines within the viral genomic RNA. Nsp15 cleavage sites (blue arrowheads) have been mapped all along the positive-sense genomic RNA and 5′-end of the negative-sense genomic RNA. Nsp15 viral RNA processing plays an important role for evading detection by the host innate immune response.

Nsp15-like endoribonucleases are a characteristic of all coronavirus family members(*3*). Biochemical experiments with recombinant Nsp15 have established that it preferentially cleaves RNA substrates 3′ of uridines, and therefore Nsp15 is commonly called endoU alluding to its cleavage specificity (*4–7*). Conservation of Nsp15 across Coronaviridae suggests that its endonuclease function is critical for their viral life cycle, however the specific role of Nsp15 in viral propagation is still unclear. Nsp15 was initially thought to play an essential proofreading role in viral replication until it was shown that the coronavirus, mouse hepatitis virus (MHV), can replicate with a catalytically-deficient variant of Nsp15 (*3, 6*). More recent work suggests that rather than functioning in viral replication, Nsp15 processes viral RNA to evade host sensors to foreign nucleic acids (*8, 9*). For instance, analysis of viral RNA from MHV-infected cells harboring a catalytically-deficient Nsp15 revealed an accumulation of 12-17 polyuridine tracts at the 5′-end of the negative-strand viral RNA intermediates. Considering that polyuridine negative-strand RNA elicits an interferon mediated response, this suggests a role for Nsp15 in regulating the length of polyuridines found at the 5′-end of negative-strand viral RNA to evade activation of the host innate immune response (Fig. 1)(*8*). High-throughput sequencing from MHV-infected macrophages also recently identified additional Nsp15 cleavage sites within the viral positive-strand RNA (Fig. 1)(*10*). Collectively, these studies suggest the existence of multiple Nsp15 cleavage targets that are important to regulate the accumulation of viral RNA and prevent activation of RNA-activated antiviral responses.

Crystal structures of Nsp15 have been reported for a number of Coronaviridae. These structures of Nsp15 have revealed a common hexameric assembly made up of a dimer of Nsp15 trimers (*4, 5, 11–13*). Each Nsp15 protomer is composed of three domains including an N-terminal domain (ND) that is important for oligomerization, a variable middle domain (MD), and a C-terminal endonuclease (endoU) domain that shares homology with other endoU enzymes (Fig. 2A)(*14*). Nsp15 is only active as a hexamer, but the molecular requirement for Nsp15 oligomerization is unknown. A crystal structure of Nsp15 from SARS-CoV-1 lacking the first 28 residues of the ND is monomeric and reveals a misfolded endoU active site suggesting that Nsp15 may rely on oligomerization as an allosteric activation switch (*12*). How Nsp15 specifically cleaves the phosphodiester bond following uridines is also poorly understood. The Nsp15 active site contains two well conserved histidine residues that are important for catalysis and reminiscent of the well characterized RNase A active site (*4, 11, 13, 15*). RNase A catalyzes a two-step reaction that first generates a 2′3′-cyclic phosphate (2′3′-cP) that is then hydrolyzed to form a 3′-phosphate (3′-P) (*16*). It is currently unclear if Nsp15 facilitates the second hydrolysis step like RNase A (*17, 18*). To gain insight into the structure and catalytic mechanism of Nsp15, we solved a series of cryo-EM reconstructions of Nsp15 from SARS-CoV-2. The cryo-EM derived atomic model identified Nsp15 residues important for uridine recognition and molecular dynamic simulations uncovered the conformational malleability of the endoU domain. Furthermore, biochemistry and mass spectrometry revealed molecular details into how the Nsp15 hexamer processes RNA.

**Fig. 2.**
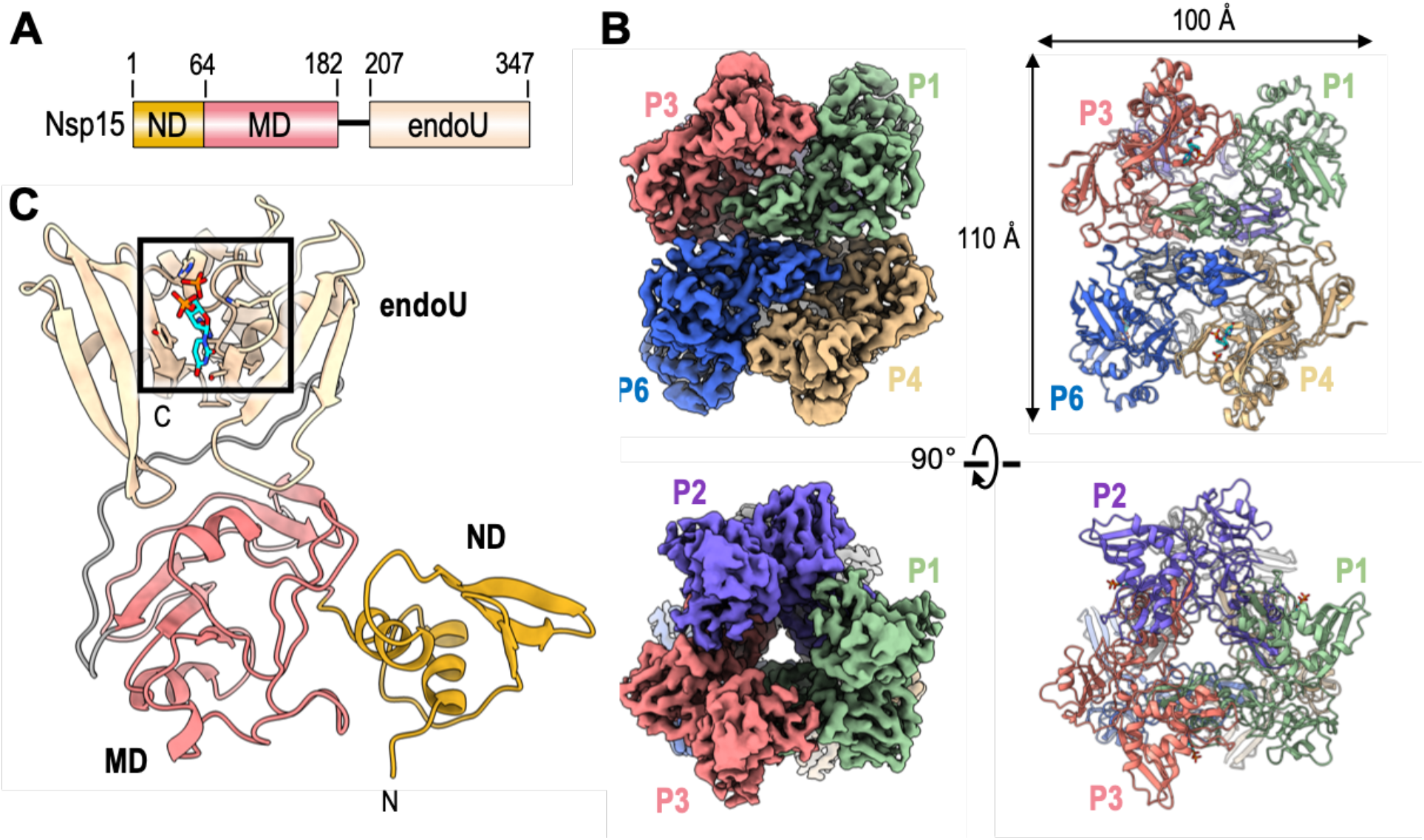
Architecture of hexameric SARS-CoV-2 Nsp15. **(A)** Domain organization of Nsp15. The numbering corresponds to the amino acid residues at the domain boundaries. The Nsp15 N-terminal domain (ND) is shown in orange, the middle domain (MD) is shown in red, and the poly-U specific endonuclease domain (endoU) is shown in beige. **(B)** Orthogonal views of the cryo-EM map reconstruction of UTP-bound Nsp15 (left) and its corresponding model shown as a cartoon (right). Each Nsp15 protomer is colored as green (P1), purple (P2), coral (P3), tan (P4), gray (P5), and blue (P6). **(C)** Model of Nsp15 protomer shown as a cartoon and colored as seen in panel A. The position of the Nsp15 endoU active site is highlighted with a black box and the modeled nucleotide is shown as sticks (cyan). N and C mark the N- and C-termini, respectively.

## Results

### Cryo-EM reconstruction reveals the hexameric arrangement of SARS-CoV-2 Nsp15

We characterized SARS-CoV-2 Nsp15 bound to a UTP nucleotide by single particle cryo electron microscopy (cryo-EM). Recombinant wild-type (wt) Nsp15 was purified as a stable hexamer using a bacterial expression system (Fig. S1). Over 1000 micrographs were collected from grids prepared with recombinant wt-Nsp15 and an excess of UTP. Following 3D classification, the particles converged into a single prominent class with a resolution of 3.36 Å (Fig. S2). The cryo-EM reconstruction of wt-Nsp15 bound to UTP revealed a hexameric assembly containing 6 protomers of Nsp15 (designated as P1-P6) (Fig. 2B) with D3 symmetry. Classification and refinement were performed with and without imposing D3 symmetry, however no asymmetric conformational states were observed. A combination of rigid-body and real space refinement was used to fit the crystal structure of SARS-CoV-2 Nsp15 (PDB ID: 6wlc) into the cryo-EM reconstruction (Table 1).

**Table 1.**
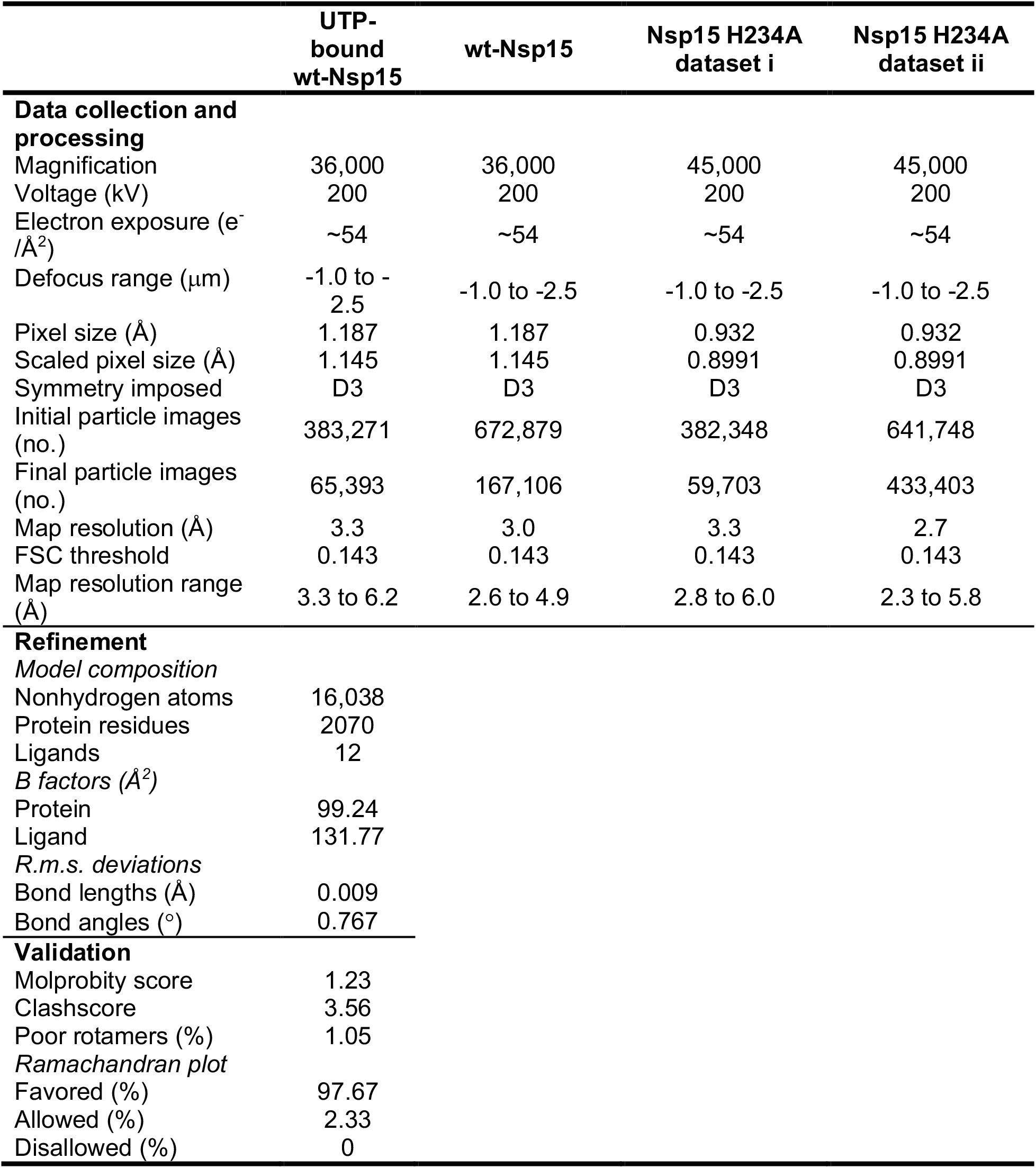
Cryo-EM data collection, refinement, and validation statistics.

For each Nsp15 molecule, all three domains of Nsp15 were clearly visible in the reconstruction as well as the uracil base and ribose sugar of the bound UTP molecule (Fig. S2). Despite adding excess manganese to the storage buffer, we did not observe discrete density for a metal ion suggesting that the stabilizing effect of manganese may be due to non-specific interactions along the highly charged patches of Nsp15 (*4, 19*). Overall the cryo-EM structure of Nsp15 is very similar to recently published crystal structures of SARS-Cov-2 Nsp15 (*11*). The barrel-shaped Nsp15 particle has dimensions of 100 × 110 Å with a narrow, negatively-charged channel that runs through the middle (Fig. 2B). The small ND domain, composed of 64 amino acid residues, mediates oligomerization of the complex by forming two head to head stacked trimers. Following the ND is the MD, composed of a mixture of beta-strands and two small alpha-helices (Fig. 2C). Following the MD is the well-conserved endoU domain containing the nuclease active site (Fig. 2C). The unique arrangement of the 6 individual Nsp15 protomers positions the endoU domains on opposite ends of the hexamer (Fig. S3).

### Cryo-EM reveals the basis for uridine specificity

The Nsp15 active site contains several residues that are well conserved amongst Nsp15 homologues and important for nuclease activity and specificity (Fig. 3A and S4). The active site within each individual Nsp15 protomer lies near the interface with a neighboring endoU domain (Fig. 3B). This close positioning of the active site to the neighboring protomer hints at a possible mechanism of allosteric communication between the Nsp15 protomers. The active site residues within the endoU domain have well defined side-chain density in the cryo-EM reconstruction along with additional density for a ligand (Fig. 3B, S2 and Mov. S1). While we used an excess of UTP when vitrifying Nsp15, we modeled uridine 5′-monophosphate (5′-UMP) into the active site because there was no observable density to account for the β- and γ-phosphates (Fig. 3C, S2). We observed additional ambiguous density next to 5′-UMP, which was modeled as a phosphate ion (Fig. 3C; model 3′-PO_4_) (Fig. S2). The ribose sugar of uridine forms van der Waals interactions with Y343, suggesting that this residue is critical for orienting the ribose within the active site for cleavage (Fig. 3C; top). Mutation of the equivalent tyrosine to alanine in SARS-CoV-1 and MERS (Middle East Respiratory Syndrome) Nsp15 leads to an almost complete loss in nuclease activity, underscoring the importance of this tyrosine residue (*4, 5*).

**Fig. 3.**
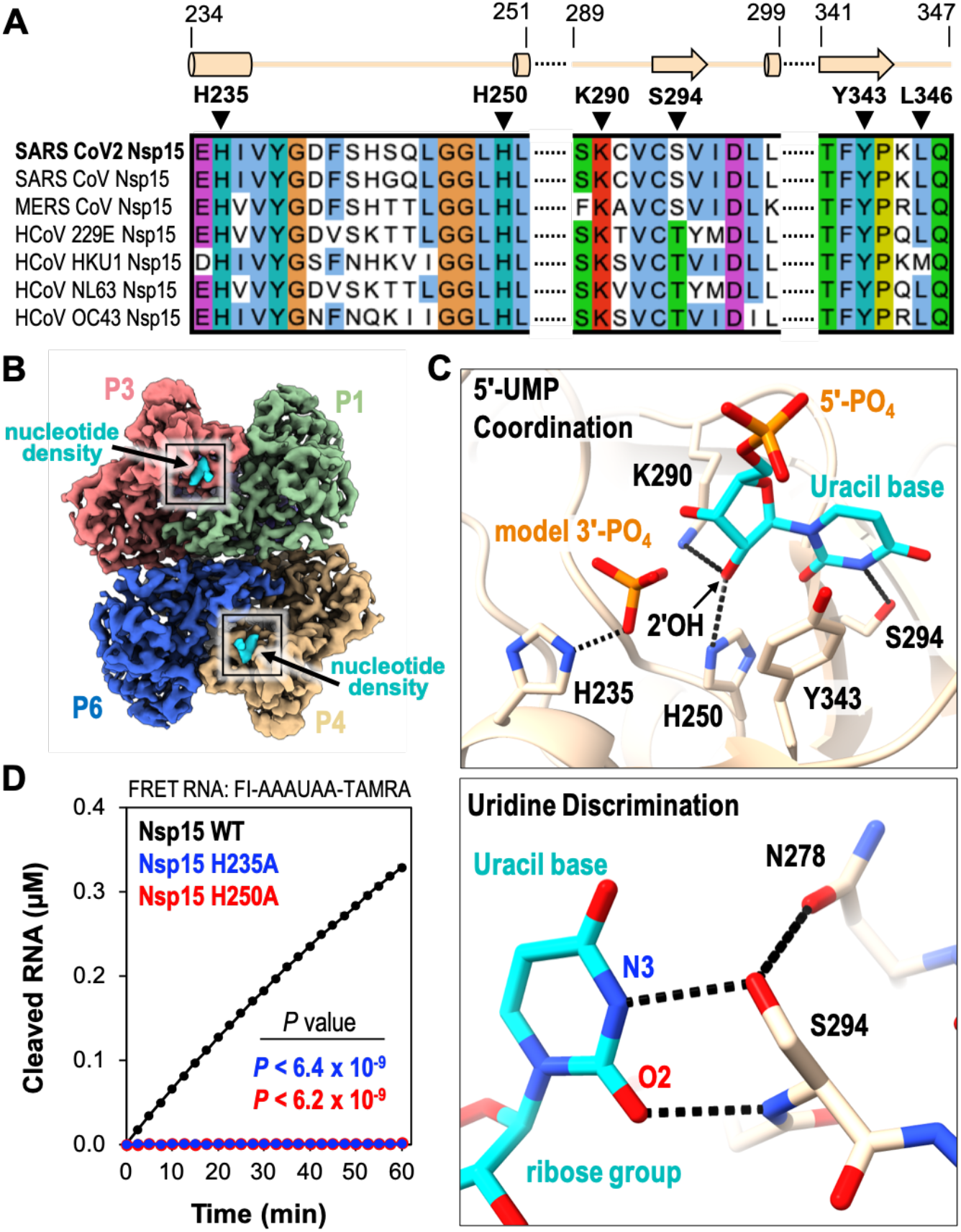
5′-UMP coordination by the Nsp15 endoU active site. **(A)** Amino acid sequence alignment of endoU active site residues from Nsp15 homologs. Secondary structure motifs observed in the nucleotide-bound Nsp15 cryo-EM structure are shown above with their corresponding amino acid residue boundaries. **(B)** Nucleotide-bound Nsp15 cryo-EM map reconstruction with protomers colored as seen in Fig. 2B. Excess UTP was added to the sample resulting in additional density within all six endoU active sites. The nucleotide density is colored in cyan and the black box demarcates the endoU active site. **(C)** Nsp15 coordination of 5′-UMP ligand. Due to the poor density of the UTP β- and γ-phosphates, 5′-UMP was modeled into the active site. Cartoon model of the 5′-UMP-bound Nsp15 endoU active site where 5′-UMP (cyan) and individual residues H235, H250, K290, S294, and Y343 are shown as sticks (top). Model of uracil base discrimination shown as sticks (bottom). Black dotted lines represent potential hydrogen bonds (2.1-3.0 Å). **(D)** RNA cleavage activity of Nsp15 variants (2.5 nM) incubated with FRET RNA substrate (0.8 μM) over time. RNA cleavage was quantified from three technical replicates. The mean and standard deviation are plotted and *p* values of wt-Nsp15 compared to H235A (blue) and H250A (red) are reported from two-tailed Student’s *t* tests.

The cryo-EM structure revealed the significance of S294 for uridine specificity. Molecular docking of 5′-UMP into the Nsp15 active site previously suggested that this serine residue may be the critical determinant for base specificity (*5*). The O2 and N3 positions of the uracil base are within hydrogen bonding distance of the S294 backbone N and side-chain OG atoms, respectively (Fig. 3C; bottom). Nsp15 N278 is also within hydrogen bonding distance of Ser294 and likely contributes to proper positioning of Ser294 within the active site. Across Nsp15 homologues the equivalent residue to S294 is either a serine or threonine (Fig. 3A). Previous work with SARS-CoV-1 Nsp15 demonstrated that mutation of this serine to a threonine does not affect activity. Yet, mutation to an alanine reduces cleavage activity by more than 5-fold and abrogates uridine specificity such that a single rU or rC containing deoxyribonucleotide are cleaved at comparable rates (*5*). While this manuscript was in preparation a series of additional Nsp15 crystal structures with different ligands were deposited in the Protein Data Bank (*20*). One of these structures includes a 5′-UMP in the active site. This crystal structure is in excellent agreement with our cryo-EM reconstruction and supports the same conclusions drawn about the role of S294 in base specificity. The crystal structure also reveals L346 is within hydrogen bonding distance with O4 of the uracil base and likely contributes to active site alignment along with uridine specificity. Interesting, L346 is not ordered in our cryo-EM reconstruction suggesting substrate binding may promote additional ordering of the active site. Thus, our cryo-EM reconstruction, recent crystal structures, and previous biochemical work support a prominent role for S294 in uracil base discrimination within the Nsp15 active site.

### Two active site histidine residues are required for nuclease activity

The active site of Nsp15 contains three residues that are required for catalysis including two histidine residues and a lysine residue, which is analogous to the active site of RNase A. Within the RNase A active site, one histidine functions as a general base to activate the 2′OH while the other histidine functions as a general acid to donate a proton to the leaving 5′OH (*16*). During the second hydrolysis step the roles of the two active site histidine residues are reversed. The three Nsp15 catalytic residues, H235, H250, and K290 are clustered around the ribose sugar of 5′-UMP (Fig. 3C; top). H250 and K290 are both within hydrogen bonding distance of the 2′OH of the ribose. The active site arrangement suggests that similar to RNase A, H250 functions as a general base to activate the 2′OH for nucleophilic attack, while K290 stabilizes the deprotonation of O2′. In contrast, H235 is set back further from the ribose and is within hydrogen bonding distance of the phosphate ion that we modeled into the active site (Fig. 3C; top). This modeled phosphate is adjacent to the 3′OH of the ribose and mimics the position of the phosphate group to be attacked by the 2′OH. Based on the active site arrangement, H235 is properly aligned to play the role of the general acid during the transesterification reaction.

To verify that our recombinant Nsp15 retained nuclease activity and to confirm the significance of the SARS-CoV-2 active site histidine residues, we adapted a FRET-based assay previously used to measure the activity of Nsp15 from MERS coronavirus (*4*). We used a short 6-mer oligonucleotide (5′-AAAUAA) that is cleaved by Nsp15 3′ to the single rU. The substrate contains 5′-fluorescein (FI) and 3′-TAMRA labels. The FI fluorescence is quenched by the TAMRA label in the uncleaved substrate, and we can monitor RNA cleavage by measuring the increase in FI fluorescence as the TAMRA label is released. We made two single mutants of Nsp15 in which we individually mutated the active site histidine residues to alanine (H235A or H250A). Both mutants purified as stable hexamers, confirming that these mutations do not disrupt the oligomerization of Nsp15 (Fig. S1). We measured RNA cleavage over a one-hour time course with wt-Nsp15 and the H235A and H250A Nsp15 variants. Wt-Nsp15 displayed robust RNA cleavage that increases as a function of time, but we could not detect any RNA cleavage with the two histidine mutants (Fig. 3D). These results confirm that recombinant Nsp15 is active and H235 and H250 are required for RNA cleavage.

### Nsp15 predominantly catalyzes 2′-*O*-transesterification

After confirming that our recombinant Nsp15 was active, we used mass spectrometry to determine the identity of the Nsp15 RNA cleavage products. First, we calculated the theoretical masses of all potential cleavage products from our 6-mer FRET RNA substrate (Table S1). Mass spectrometry can identify the site of RNA cleavage. Moreover, mass spectrometry can distinguish the mass difference of 18.01 Da for the two possible 3′-end products (the 2′3′-cP intermediate versus the 3′- P hydrolyzed intermediate). We analyzed the 6-mer FRET RNA substrate using liquid chromatography electrospray ionization mass spectrometry (LC-ESI-MS) in the absence of Nsp15 and observed a MS spectra peak of 3294.74 Da, corresponding to the theoretical molecular mass of the uncleaved RNA substrate (Fig. S5A and Table S1). Next, we analyzed the RNA products following a 30-minute enzymatic reaction with Nsp15. Similar to other Nsp15 homologs, we observed products resulting from a single cleavage site 3′ to the uridine (5′-AAAU^AA), confirming Nsp15’s strict specificity for uridine. We observed a prominent peak in the extracted ion chromatogram with a calculated mass of ~1463 Da, which was absent in our negative control (Fig. 4A). The experimental mass of this RNA product corresponds well with the theoretical mass of 5′-HO-AA-TAMRA (Table S1). The MS spectrum of the 5′-HO-AA-TAMRA product has a mass of 1464.43 Da and in conjunction with MSMS data confirms its identity (Fig. 4B and S5B). Furthermore, we could not detect any other 5′ cleavage products, confirming Nsp15 exclusively cuts the RNA substrate 3′ of the uridine. A second prominent peak was also detected in the extracted ion chromatogram that was absent in our negative control and corresponds to a 3′ RNA product (Fig. 4C). The MS spectrum of the 3′ product revealed a doubly charged ion at m/z 914.14 and, hence, a molecular mass of 1830.30 Da corresponding to 5′-FI-AAAU-2′3′-cP and another doubly charged ion at m/z/ 923.14 and, hence, a molecular mass of 1848.31 Da corresponding to 5′-FI-AAAU-3′-P (Fig. 4D, S5C and Table S1). To determine which 3′ RNA product is more prominent, we normalized and compared the abundance of the 2′3′-cP and 3′-P products assuming similar ionization efficiencies of the 2′3′-cP and 3′-P products (Fig. 3E). We observed that the 2′3′- cP makes up 80% of the total 3′-product confirming the 2′3′-cP is the major cleavage product and suggesting the rate of hydrolysis is slow.

**Fig. 4.**
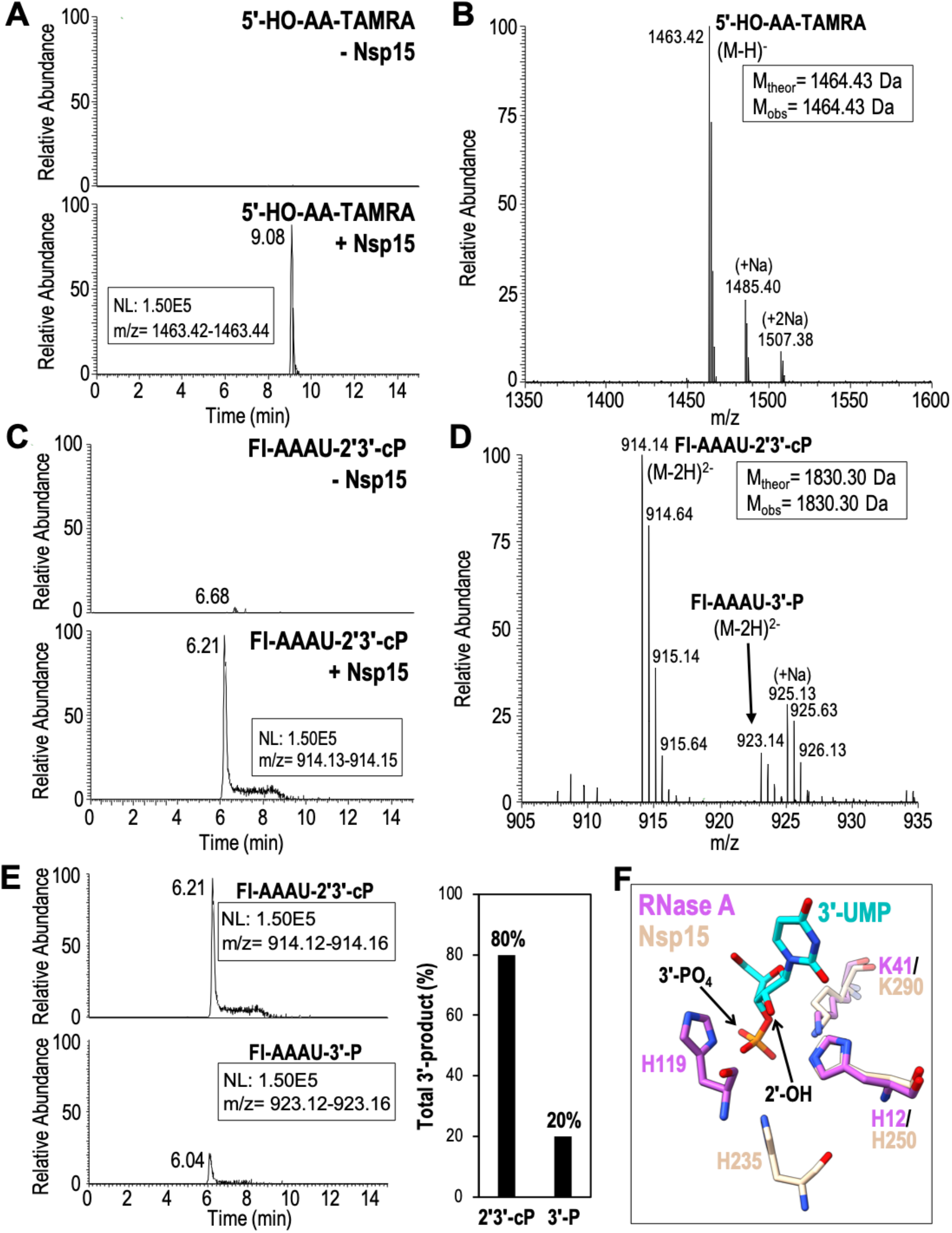
Extracted ion chromatograms and MS spectra of Nsp15 RNA cleavage products. **(A)** Extracted ion chromatograms of the 3′-TAMRA labeled AA-TAMRA RNA cleavage product. 5′- HO-AA-TAMRA is readily observed in the presence of Nsp15 (+Nsp15) as a singly charged ion at m/z 1463.42, but is undetectable in the absence of enzyme (-Nsp15). **(B)** MS spectrum confirms the identity of 5′-HO-AA-TAMRA and does not detect the presence of alternative 5′-cleavage products. **(C)** Extracted ion chromatograms of the 5′-fluorescein labeled (FI) FI-AAAU cleavage product. FI-AAAU is only detected in the presence of Nsp15. **(D)** MS spectrum confirms the identity of FI-AAAU. The 3′-product is observed primarily as a doubly charged ion at m/z 914.14, which corresponds to a 2′3′-cP terminated moiety. A doubly charged ion at m/z 923.14 is also present and corresponds to a 3′-P terminated species. MSMS spectra of m/z 1463.14 and m/z 914.14 unambiguously confirm the identity of these ions (Fig. S5) **(E)** Extracted ion chromatograms of the 5′-FI-AAAU-2′3′-cP cleavage product (top) and the 5′-FI-AAAU-3′-P cleavage product (bottom) set to the same scale to demonstrate that under the conditions employed, the majority of the cleavage product is terminated by a cyclic phosphate. The graphical representation of the areas under the curve of the extracted ion chromatograms show that approximately 80% of the cleavage product is the cyclic phosphate (assuming similar ionization efficiencies of the two species). **(F)** Superimposition of SARS-CoV-2 Nsp15 (beige) and *Bos taurus* RNase A (PDB ID: 1o0n; magenta) active site residues bound to 3′-UMP (cyan) shown as sticks.

We compared the active sites of Nsp15 and RNase A to determine why Nsp15 does not promote the hydrolysis step as efficiently as RNase A. We superimposed the active sites of Nsp15 and RNase A using a crystal structure of RNase A solved in the presence of 3′-UMP (*21*). H250 and K290 of Nsp15 superimpose well with the equivalent H12 and K41 residues from RNase A (Fig. 4F). In contrast, H235 from Nsp15 and H119 from RNase A do not superimpose (Fig. 3F). H119 of RNase A is near the ribose group and is fixed into position by a network of hydrogen bonds and tightly coupled to a carboxyl (Asp121) to promote catalytic action. Conversely, H235 of Nsp15 is ~ 8 Å away from the ribose group and therefore it is not positioned to promote hydrolysis of the cyclic phosphate. This difference in Nsp15 active site architecture likely slows the hydrolysis reaction leading to an accumulation of 2′3′-cP products. This is reminiscent of the active site of a related nuclease Angiogenin (*22*). The active site of Angiogenin is partially blocked by a glutamine residue that is not found in RNase A (*23*). The glutamine residue alters the active site geometry in such a way that cleavage is not as efficient as RNase A and it primarily produces 2′3′-cP products.

### UTP binding stabilizes endoU domain conformation

In addition to determining the structure of Nsp15 in the presence of UTP, we also determined several cryo-EM reconstructions of Nsp15 in the absence of ligand at resolutions ranging from 2.9 to 3.3 Å (Fig S6-S8). First, we vitrified the H235A-Nsp15 mutant, which cannot cleave RNA, in the presence of a 21-nucleotide single-strand RNA (SS-RNA) substrate in an attempt to solve the structure of Nsp15 bound to RNA. We processed this data with and without imposing D3 symmetry, but did not observe any asymmetric states or the presence of RNA. Overall the cryo-EM reconstruction of apo H235A-Nsp15 are very similar to the uridine-bound reconstruction with one notable difference, an observed loss in the local resolution of the endoU domain (Fig. S6). One possibility for this loss in resolution is the presence of heterogeneity in the conformation of the endoU domain. We carried out 3D variability analysis to model the conformational landscape of the endoU domain present in our cryo-EM data (*24*). This analysis revealed multiple conformational states of the endoU domain with respect to the remaining Nsp15 hexamer (Fig. 5A and Mov. S2). The conformational variability of the endoU domain is restrained, but suggests that in the absence of a ligand the endoU domain wobbles towards the central channel with respect to the ND and MD domains and is not locked into a fixed position.

**Fig. 5.**
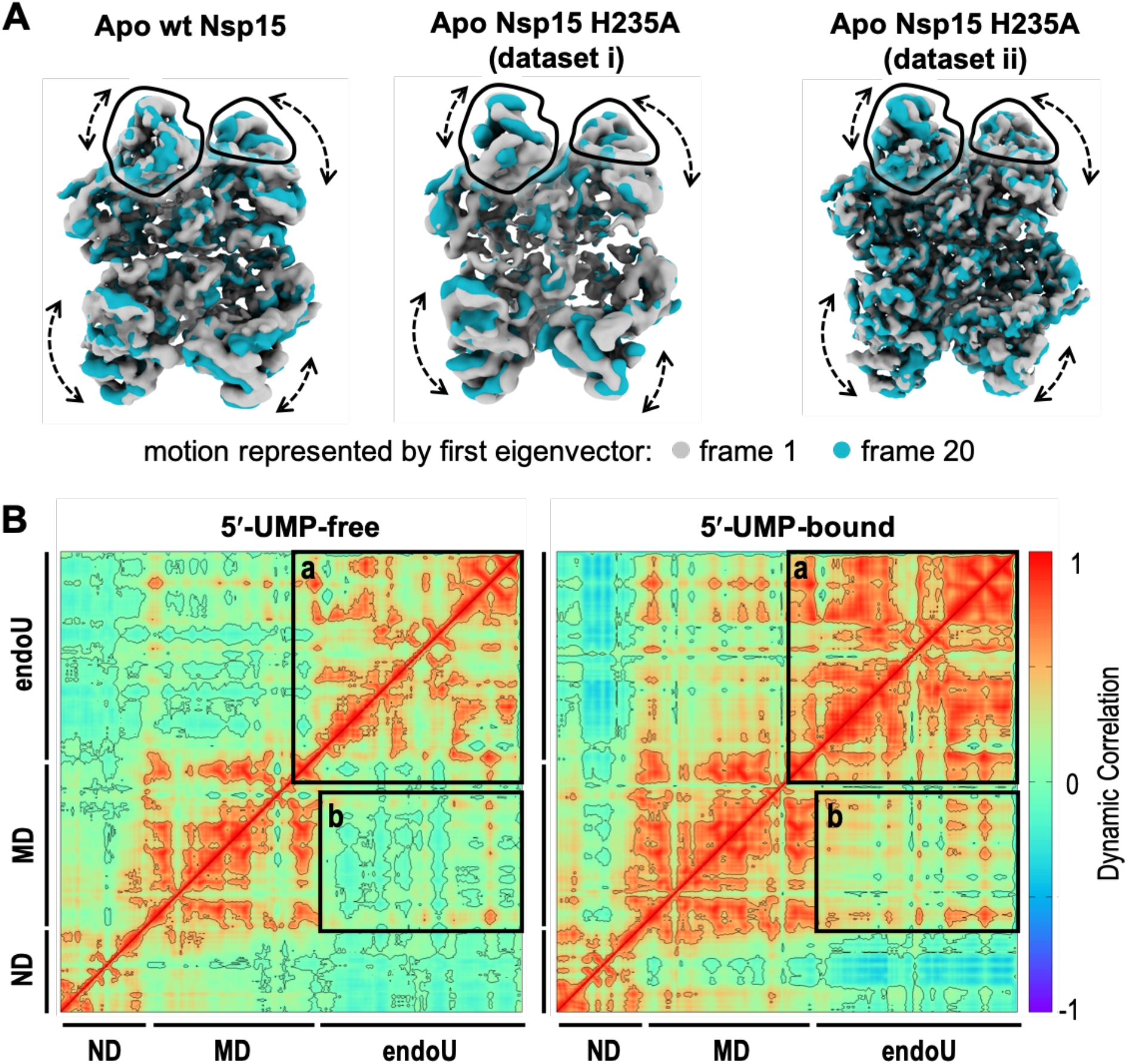
Conformational heterogeneity of the Nsp15 endoU domain. **(A)** 3D variability analysis (*30*) of cryo-EM reconstructions of Nsp15 apo-states. Frame 1 (gray) and frame 20 (blue) of a 20-frame set describing the motion by the first eigenvector of the cryo-EM reconstructions. Black outlines demarcate the unique views of the endoU domains and dotted arrows demarcate regions of conformational heterogeneity. **(B)** Dynamical cross-correlation matrix (DCCM) analysis of a 5′-UMP-bound protomer and its 5′-UMP-free form from hexamer systems. Both inter- and intra-domain positive correlations are significantly enhanced due to 5′-UMP-binding. The correlations were calculated from the equally spaced 100-configurations extracted from the last 500 ns of each simulation. Black boxes highlight the positive correlated motions between the endoU domains within the hexameric assembly (box a) and the endoU domain relative to the N-terminal half of the protomer (box b).

In order to confirm that the observed wobbling of the endoU domain was not due to the H235A active site mutation or the presence of RNA when vitrifying the sample, we determined two additional cryo-EM structures of wt-Nsp15 and Nsp15 H235A vitrified in the absence of any ligand. Both of these reconstructions also display a loss in the local resolution of the endoU domain in comparison with UTP-bound Nsp15 (Figs. S7-S8). We repeated 3D variability analysis on these datasets and observed similar conformational landscapes of the endoU domain (Fig. 5A and Movies S3-S4). Collectively the three apo-Nsp15 cryo-EM reconstructions reveal that the endoU domain of Nsp15 is dynamic and samples multiple conformations in its ligand-free state.

Finally, we used molecular dynamics simulations to confirm the observed wobbling of the endoU domain in the absence of ligand. We looked at the dynamics of the Nsp15 monomer and Nsp15 hexamer in the presence and absence of 5′-UMP. We established the stability of the Nsp15 monomer and hexamer by determining the root mean square deviations (RMSD) from molecular dynamics trajectories following the convergence of the RMSD values in the last 100 ns of the simulations (Fig. S9). There is a significant difference in the RMSDs of the monomer compared to the averaged values of individual monomers from the hexamer both in the absence (2.0 vs 1.3 Å) and presence (2.3 vs 1.4 Å) of 5′-UMP. Thus, the hexamer provides stability for the individual Nsp15 protomers and this stability likely contributes to the requirement of Nsp15 oligomerization for RNA cleavage activity.

Next, we determined the impact of 5′-UMP binding on the dynamics of the monomer and hexamer. Binding of 5′-UMP does not result in large changes in the thermal fluctuations, but results in differences with the dynamic correlation (Fig. S9). Dynamic correlation reflects the association between the distances of the alpha-carbons across the entire protein backbone. In the absence of 5′-UMP we observe correlations within the residues of individual domains of Nsp15, but there is little correlation among the three distinct domains. This suggests the ND, MD, and endoU domains are largely moving independently of one another in the absence of a ligand. In contrast, we observe an increase in the correlated motions of both the residues within the endoU domains across the hexamer and among the ND, MD, and endoU domains in the presence of 5′-UMP (Fig. 5B and Table S2). This observation is in-line with 5′-UMP-induced ordering of the Nsp15 endoU domain and a previous report that the addition of mononucleotide and dinucleotide analogs significantly increase Nsp15 stability by differential scanning fluorometry (*20*). Thus, 5′-UMP binding locks the endoU domain into a fixed position that correlates its motion with the rest of the Nsp15 higher-order assembly.

## Discussion

Here we present a cryo-EM derived atomic model of the hexameric endonuclease Nsp15 from SARS-CoV-2. In the presence of 5′-UMP, Nsp15 has an ordered active site that draws many parallels with the well-studied RNase A, which utilizes an elegant acid-base reaction mechanism to catalyze a two-step reaction of transesterification and hydrolysis. Mass spectrometry revealed that analogous to RNase A, Nsp15 can catalyze both reaction steps, however we observed a significant accumulation of the 2′3′-cP product from the transesterification reaction indicating that hydrolysis is slow. Based on the comparison of the active sites of Nsp15 and RNase A this difference can be attributed to an alternative location of H235 in the Nsp15 active site. While the nature of the 3′-end of the cleavage product can have important implications for downstream RNA processing and/or function (*22*), it seems unlikely that the identity of the 3′-end is critical for viral pathogenesis. Nsp15, with a presumptive role in regulating the length of poly(U) tails at the 5′-end of the negative-strand (Fig. 1), generates a shortened negative-strand with a 5′-hydroxyl and a range of short U-ended oligonucleotide products with a mixture of 2′3′-cP and 3′-P ends (*8*).

The 5′-UMP bound cryo-EM structure is similar to existing Nsp15 crystal structures, however, the apo cryo-EM reconstructions revealed structural heterogeneity within the endoU nuclease domain. This heterogeneity demonstrates that the endoU domain wobbles with respect to the N-terminal half of Nsp15 in the absence of RNA ligand. Molecular dynamics confirmed that in the absence of ligand the endoU domain moves independent of the rest of the molecule. In contrast, in the presence of 5′-UMP the endoU domain is locked down and its movement is correlated with the N-terminal half of the protein. While more work is needed to establish the functional significance of the endoU wobble, there are several possibilities as to why this could be critical for Nsp15 function. Flexible active sites that become stiff upon engaging substrates have been shown to promote catalysis by reducing the substrate-binding energy (*25*). Another possibility is that the wobbling of the endoU domain is important for Nsp15 to accommodate different RNA substrates. Recent work suggests that in addition to cleaving the 5′-end of the negative-strand, Nsp15 can cleave multiple sites within the viral RNA to prevent activation of host anti-viral sensors (*8, 10*). Finally, wobbling of the endoU domain may provide Nsp15 with a mechanism of allosteric communication between the six individual protomers. While we only observed Nsp15 particles with six-fold symmetry, we hypothesize that binding of large RNA substrates will lead to the formation of asymmetric states that rely on coordination between the protomers to bind and process RNA. This is further supported by the free energy estimations from molecular dynamics simulations, which revealed that 5′-UMP initially binds all 6 active sites, but only remains stably associated with a single protomer (Table S2).

In summary, our cryo-EM reconstructions, biochemistry, and molecular dynamics reveal critical insight into how SARS-CoV-2 Nsp15 processes viral RNA. The series of structures presented in this manuscript can be used to aid in the development of effective inhibitors which are urgently needed as reported cases of Covid-19 continue to rise. Nucleotide analogues have shown great promise as viral inhibitors against SARS-CoV-2, thus uridine derivatives that could bind the Nsp15 active site should continue to be explored as new therapeutic targets (*20*). Moreover, recent in silico-based approaches have identified potential Nsp15 inhibitors that await structural and biochemical validation (*26, 27*).

## Materials and Methods

### Nsp15 construct design

The nucleotide sequence for the Tobacco Etch Virus (TEV) recognition site was added upstream of full-length SARS-CoV-2 *Nsp15* that was codon optimized for *E. coli* expression. The TEV-Nsp15 construct was inserted into the bacterial expression vector pET-14b in frame with the N-terminal 6x His tag and Thrombin cleavage site to create the 6X-His-Thrombin-TEV wt-Nsp15 construct. Nsp15 catalytic-deficient variants H235A and H250A were created using the wt-Nsp15 template. All plasmids were verified by DNA sequencing (GeneWiz).

### Protein expression of Nsp15 variants

Wt-Nsp15 was overproduced in *E. coli* CD41 (DE3) competent cells in Terrific Broth supplemented with 100 μg/mL ampicillin. Transformed cell cultures were grown to an optical density (600 nm) of 1.0 prior to their induction with 0.2 mM Isopropyl β-D-1-thiogalactopyranoside (IPTG) and incubated for 3 hours at 37°C. Harvested cells were stored at - 80°C until needed. Catalytic-deficient Nsp15 variants (H235A or H250A) were overexpressed in *E. coli* Rosetta (DE3) pLacI competent cells in Terrific Broth supplemented with 100 μg/mL ampicillin and 25 μg/mL chloramphenicol. Transformed cell cultures were grown to an optical density (600 nm) of 1.0. Cell cultures were stored at 4°C for 1 hour prior to their induction with 0.2 mM IPTG and overnight incubation at 16°C. Harvested cells were stored at −80°C until needed.

### Protein purification

Cells were resuspended in Lysis Buffer (50 mM Tris pH 8.0, 500 mM NaCl, 5% glycerol, 5 mM β-ME, 5 mM imidazole) supplemented with cOmplete EDTA-free protease inhibitor tablets (Roche) and disrupted by sonication. The lysate was clarified at 26,915 × g for 50 minutes at 4°C and then incubated with TALON metal affinity resin (Clontech). His-Thrombin-TEV-Nsp15 variants were eluted with 250 mM imidazole and further purified by gel filtration on a Superdex-200 column (GE Healthcare) in SEC Buffer (20 mM Hepes pH 7.5, 150 mM NaCl, 5 mM MnCl_2_, 5 mM β-ME). To remove the N-terminal 6x His-tag from His-Thrombin-TEV-Nsp15, variants were incubated with thrombin (Sigma) at room temperature in Thrombin Cleavage Buffer (50 mM Tris pH 8.0, 150 mM NaCl, 5% glycerol, 2 mM β-ME, 2 mM CaCl_2_) for 3 hours. Thrombin cleavage was quenched by the addition of 1 mM PMSF (phenylmethylsulfonyl fluoride). Proteolytic cleavage reactions were incubated with TALON metal affinity resin and tagless protein was eluted in batch and resolved by gel filtration using a Superdex-200 column equilibrated with SEC Buffer.

### Preparation of Nsp15 cryo-EM specimens

Purified wt-Nsp15 and catalytic-deficient Nsp15 H235A were resolved over a Superdex-200 (GE Healthcare) gel filtration column using SEC Buffer to isolate the hexamer. Prior to grid preparation Nsp15 variants were diluted into low-salt buffer (20 mM Hepes pH 7.5, 100 mM NaCl, 5 mM MnCl_2_, 5 mM β-ME). Nsp15 variants (0.75 μM) were incubated in the presence of excess 8 μM 22-nucleotide uridine-rich single-stranded RNA (UUUAGGUUUUACCAACUGCGGC/36-TAMSp/) (H235A-APO, dataset i), in the absence of ligand (H235A-APO, dataset ii), in the presence of excess 333 μM 22-nucleotide single-stranded RNA (SS-RNA: 5′- ACGUACGCGGAAUACUUCGAA-TAMRA-3′) (wt-APO), and in the presence of excess 2 mM uridine 5′-triphosphate (wt-UTP bound) for 1 hour at 4°C. UltrAuFoil R1.2/1.3 300 mesh gold grids (Quantifoil) were rendered hydrophilic using the Tergeo plasma cleaner (Pie Scientific). Protein mixture (5 μL) were deposited onto the grids and blotted for 3 seconds using an Automatic Plunge Freezer (Leica).

### Data acquisition and image processing

Images of Nsp15 variants were collected on a Talos Arctica electron microscope (Thermo Fischer Scientific) operated at 200 keV and equipped with a K2 Summit direct detection camera (Gatan). Movies were recorded in counting mode at a nominal magnification of 45,000x or 36,000x corresponding to 0.932 Å/pixel and 1.187 Å/pixel respectively (see Table 1). Total exposure time was 8.4 seconds at a dose rate of 6.54 e^−^Å^−2^s^−1^ resulting in a total dose of ~54 e^−^A^−2^, distributed over 60 frames. Images were recorded across a defocus range of −1.0 to −2.5 μm.

Beam-induced motion and drift were corrected using MotionCor2 (*28*) and aligned dose-weighted images were used to calculate CTF parameters using CTFFIND4 (*29*). CryoSPARC v2 (*30*) was used in all subsequent image processing. Particles were selected by template-based particle picking, downsampled by a factor of 4, extracted with a box size of 64 and subjected to an initial round of 2D classification. Full resolution particle projections from “good” classes were re-extracted using a box size of 256. Ab initio reconstruction with 3 classes was used to generate initial models. Three independent 3D refinement cycles were performed while applying C1, C3, and D3 symmetry respectively. Visual examination of the refined maps did not revealed significant differences between asymmetric units. For this reason, refined maps with D3 symmetry were used for all subsequent model building and analysis. Local resolution was calculated using cryoSPARC’s own implementation. Particles from each refinement were then post-processed using per-particle motion correction and global CTF refinement before undergoing a subsequent iteration of non-uniform refinement. Local refinement of the asymmetric unit of the map was performed by masking the asymmetric unit. The mask was padded by 5 pixels with a soft mask extended by 3 pixels. Each dataset was D3 symmetry expanded and locally refined with the fulcrum point defined as the center of the mask. 3D variability analysis was performed using cryoSPARC’s own implementation (*24*). The full mask from 3D refinement of each respective dataset was used, along with a D3 symmetry expanded dataset. A 5 Å filter resolution and 20 Å high pass resolution were used to capture movements of each asymmetric unit. Maps were visualized using the simple 3D variability display job with a total of 20 frames. Superposition of the model versus high resolution crystal structures revealed an error in the voxel size. Therefore the map was rescaled by 0.9647, to optimize the RMSD fit of the N-terminal domain to crystal structures PDB ID 6WLC and 6X4I.

### Model building

The SARS-CoV-2 Nsp15 crystal structure bound to 5′-UMP (PDB ID: 6wlc) was fit into the UTP-bound cryo-EM reconstruction using rigid-body docking in Phenix (*31*). The nucleotide density for the uracil base, ribose sugar, and α-phosphate was visible, however the density for the β- and γ-phosphates of the UTP molecules was ambiguous. For this reason, we modeled 5′-UMP. Additional density was also observed adjacent to the nucleotide within the active site. We modeled phosphate into this density, however the identity remains unknown. Fit was improved using a combination of rigid body and real-space refinement in Phenix (*31*) combined with iterative rounds of building in COOT (*32*). Hydrogen bond restraints were included for the UMP ligand to stabilize refinement in the poorly resolved ligand density. Molprobity (*33*) was used to evaluate the model and the model statistics are listed in Table 1. Figures and videos were prepared with PyMOL, Chimera, and ChimeraX (*34–36*). Model building for the apo-state cryo-EM reconstructions were not performed due to poor endoU domain density caused by conformational heterogeneity.

### Nsp15 endoribonuclease assay

Real-time Nsp15 RNA cleavage was monitored as previously described (*4*) with minor modifications. The 5′-fluorescein (FI) label fused to the FRET RNA substrate is quenched by its 3′-TAMRA label (5′-FI-AAAUAA-TAMRA-3′). The FRET RNA substrate (0.8 μM) was incubated with a constant amount of tagless Nsp15 variant (2.5 nM) in RNA Cleavage Buffer (20 mM Hepes pH 7.5, 75 mM NaCl, 5 mM MnCl_2_, 5 mM β-ME) at 25°C for 60 minutes. RNA cleavage was measured as an increase in fluorescein fluorescence. Fluorescence was measured every 2.5 minutes using a POLARstar Omega plate reader (BMG Labtech) set to excitation and emission wavelengths of 485 ± 12 nm and 520 nm, respectively. Three technical replicates were performed to calculate the mean, standard deviation, and two-tailed Student’s *t* tests.

### Mass spectrometry analysis of RNA cleavage products

To analyze the masses of the RNA cleavage products produced by Nsp15, the FRET RNA substrate (0.8 μM) was incubated in the absence and presence of tagless Nsp15 variant (2.5 nM) in RNA Cleavage Buffer at 25°C for 30 minutes. Mass spectrometry was performed essentially as described in (*37*) with the following modifications. Buffer A was 400 mM hexafluoro-2-propanol, 3 mM triethylamine (pH 7.0) and buffer B was methanol. Additionally, parallel reaction monitoring (PRM) analyses were included in the MS analyses with included masses of m/z 914.14; 923.14; 1463.42.

### Molecular dynamics

Initial monomer structures were selected from the crystal structure of the SARS-CoV-2 Nsp15 endoribonuclease bound to 5′-UMP (PDB ID: 6wlc). Four simulations systems were constructed, (a) a ligand-free monomer, (b) a 5′-UMP-bound monomer, (c) a ligand-free hexamer, and (d) a 5′- UMP-bound hexamer (with six 5′-UMP molecules). After introducing protons, each structure was solvated in a box of water; counter ions were added; and additional Na^+^ and Cl^−^ ions were placed so that the salt concentration was 100 mM. The closest box boundary is at least 15 Å away from any protein atom. The charges for 5′-UMP were generated at the 6-31g*/B3LYP level with the FF14SB force field for amino acids. After proper equilibration of each system over 30 ns under various conditions, the CUDA implementation of the Pmemd module of Amber.18 (*38*) was used to simulate unconstrained dynamics for 500 ns for monomer systems at 2 fs time step and 300K under constant pressure. The hexamer trajectories were extended over 800 ns. The MMGBSA module of Amber.18 was implemented in free energy estimations.

## Supplementary Materials

This article includes the following supplemental materials:

Figures S1 to S9

Tables S1 to S2

Movies S1 to S4

## Acknowledgements

We thank Drs. Traci Hall and Joseph Rodriguez for their critical reading of this manuscript. We would like to thank all the members of the Molecular Microscopy Consortium at the NIEHS for their help with cryo-EM data collection and processing, along with all of the members of the Mass Spectrometry Research and Support Group at the NIEHS for their help with mass spectrometry data collection and analysis. We are also grateful to Dr. Oliver Clarke for his support with cryo-EM data processing using cryoSPARC.

## Author Contributions

R.E.S. and M.C.P conceived and designed the study. M.C.P, L.C.B, J.M.K. S.K., C.K.H., J.G, and Z.D.S performed cryo-EM analysis and structure determination and refinement. L.C.B, A.H., V.D., and M.B. collected and processed cryo-EM data. M.N.F. purified all recombinant proteins. S.K expressed recombinant proteins. C.K.H and M.S. generated recombinant DNA and other materials. J.G.W. and L.J.D. processed and analyzed mass spectrometry data. L.P. performed molecular dynamics stimulations. M.C.P, L.B.D, M.L.F, C.K.H, J.G., J.G.W., L.J.D, and R.E.S. prepared the figures. M.C.P and R.E.S. wrote and revised the manuscript.

