## Supplemental Information for "Cryo-EM Structures of the SARS-CoV-2 Endoribonuclease Nsp15"

**This PDF file includes:**

Figures S1 to S9
Tables S1 to S2
Legends for Movies S1 to S4

**Other Supplementary Material for this manuscript includes the following:**

Movies S1 to S4

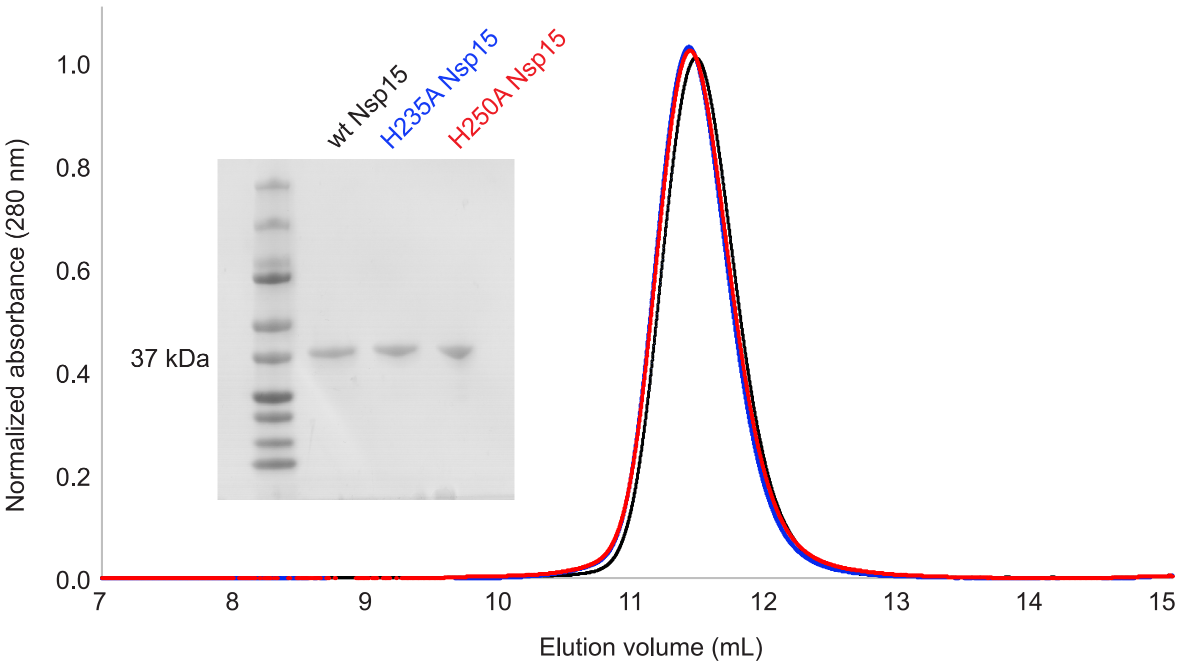

**Fig. S1. Purification of recombinant SARS-CoV-2 Nsp15 variants.** Purified Nsp15 variants wild-type (black), H235A (blue), and H250A (red) were resolved over a Superdex-200 increase 10/300 GL gel filtration column using SEC buffer. Inset is an SDS-PAGE analysis of recombinant Nsp15 wild-type (wt), H235A, and H250A variants.

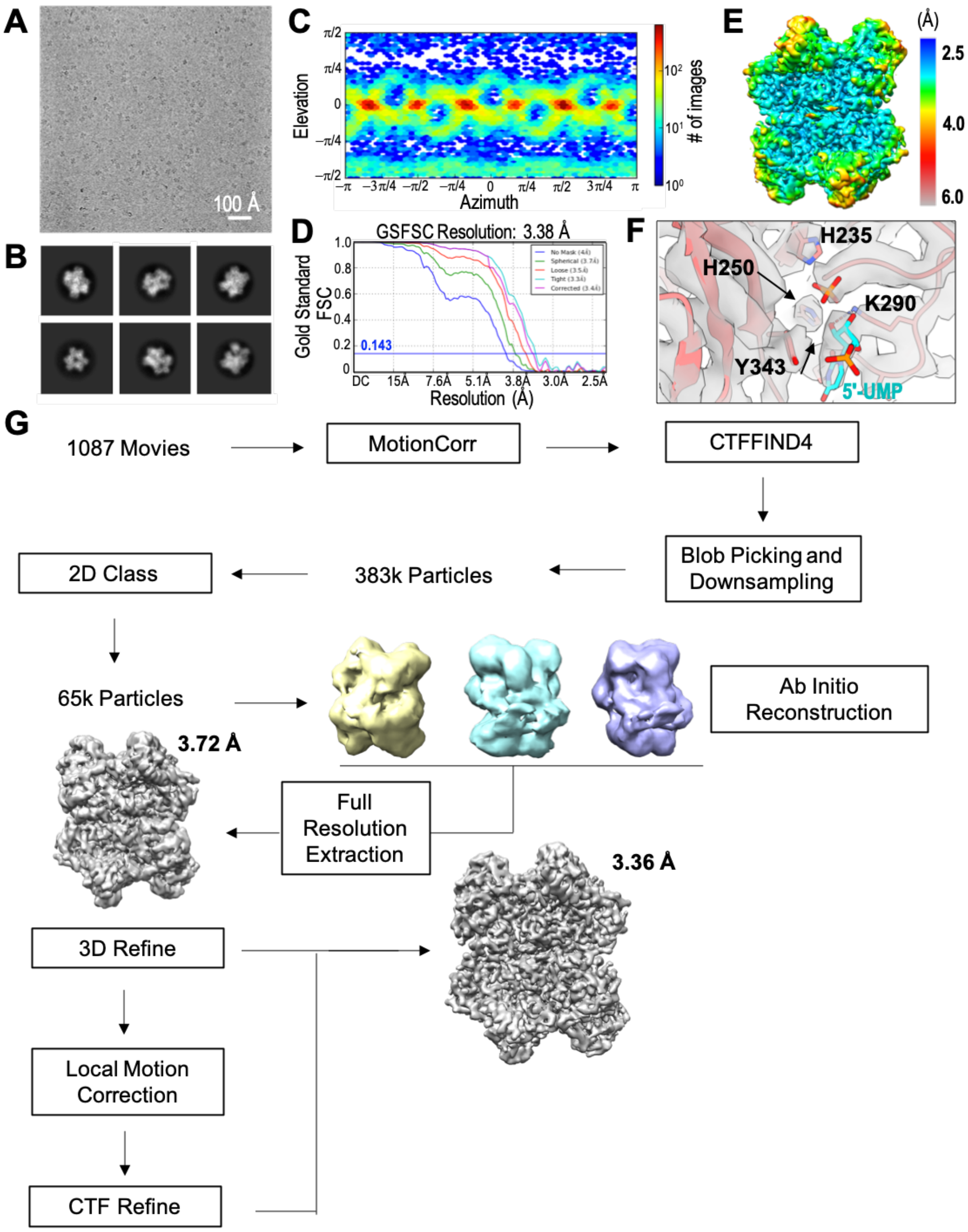

**Fig. S2. Overview of cryo-EM processing scheme for UTP-bound Nsp15. (A)** A representative micrograph of wt-Nsp15 in the presence of excess UTP in vitreous ice and **(B)** selected 2D classes generated from 1087 movies collected from an UltrAuFoil R1.2/1.3 300 mesh grid. **(C)** Angular distribution of UTP-bound wt-Nsp15 particles. **(D)** Fourier shell correlation (FSC) curve for the UTP-bound Nsp15 reconstruction. The overall resolution is 3.38 Å according to the FSC 0.143 criteria (*1, 2*). **(E)** Cryo-EM reconstruction of UTP-bound Nsp15 colored based on local resolution calculated using cryoSparc v2 (*3*). **(F)** Cryo-EM density for Nsp15 endoU active site (gray) with the corresponding model shown as a cartoon with individual active site residues, 5′-UMP, and modeled phosphate shown as sticks. **(G)** Cryo-EM processing workflow. Picked particles (383,271) were subjected to 2D classification, 3D classification, and refinement prior to local refinement of a single Nsp15 protomer in cryoSPARC v2 (*3*).

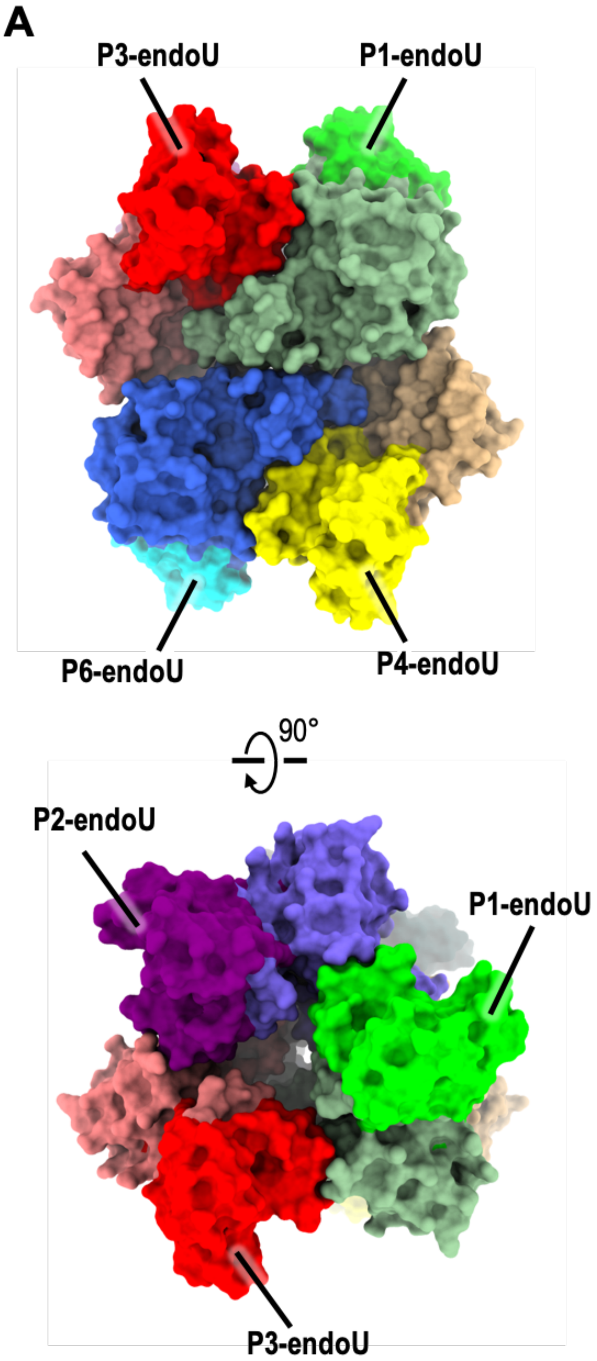

**Fig. S3. Arrangement of Nsp15 endoU domains.** Orthogonal views of **(A)** surface rendering and **(B)** cartoon model of hexameric Nsp15. The Nsp15 ND and MD domains are colored as seen in Fig. 2B. The endoU domains of six Nsp15 protomers (P1 to P6) are colored in bright hues of green, purple, red, yellow, gray, and cyan, respectfully.

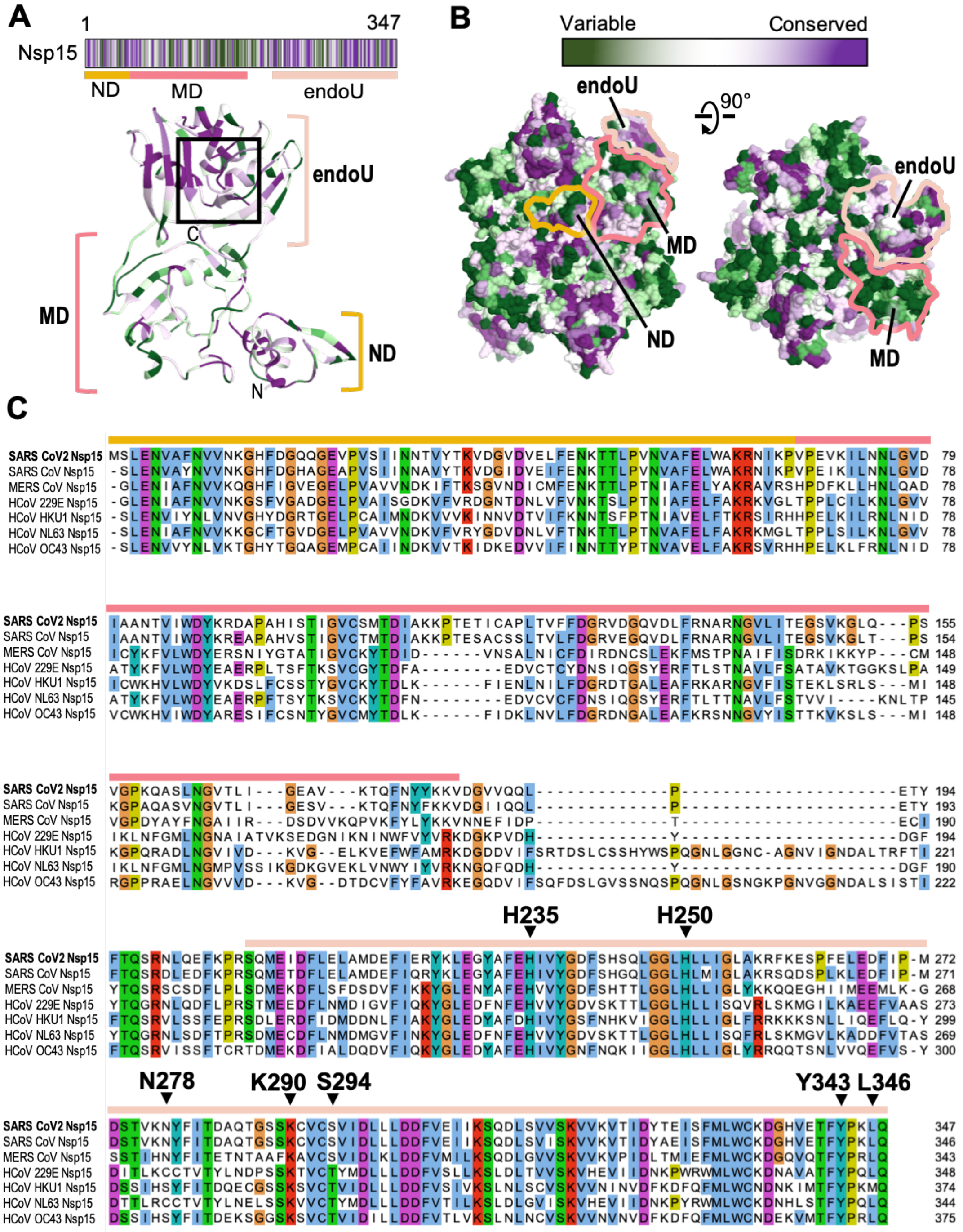

**Fig. S4. Sequence alignment and conservation of Nsp15. (A)** Consurf analysis (http:// consurf-hssp.tau.ac.il) of Nsp15 homologues plotted with respect to its domain architecture. Conserved and variable amino acid residues are colored as purple and green bars, respectively. The SARS-CoV-2 Nsp15 model of the protomer is shown as a cartoon and colored based on amino acid residue conservation. **(B)** Surface representation of hexameric Nsp15 illustrating conserved (purple) to variable (green) residues based on the Consurf analysis. The ND, MD, and endoU domains are outlined in orange, red, and beige, respectfully. **(C)** Sequence alignment of Nsp15 performed by PROMAL3D (*4*) and illustrated by JalView (*5*). Black arrowheads mark the position of Nsp15 endoU residues located within the active site.

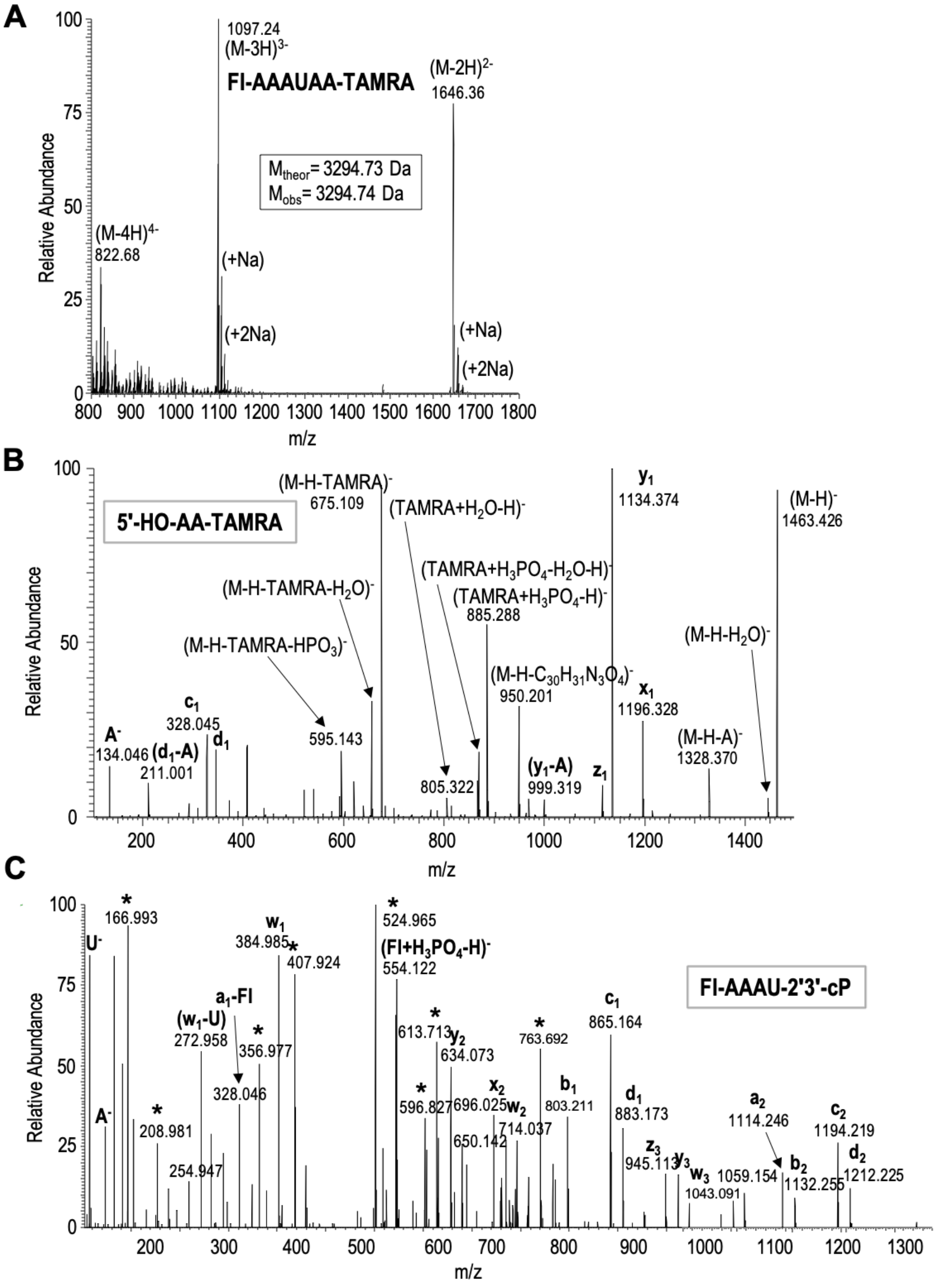

**Fig. S5. RNA analysis using mass spectrometry. (A)** The MS spectrum of the uncut FI-AAAUAA-TAMRA synthetic oligonucleotide shows doubly and triply charged species that correspond in mass to the theoretical mass of the intact oligonucleotide. **(B)** The MSMS spectrum of the m/z 1463.42 ion yields extensive fragmentation (*6*) that confirms the identity of 5′-HO-AA-TAMRA cleavage product. **(C)** The MSMS spectrum of the m/z 914.14 ion yields extensive fragmentation (*6*) that confirms the identity of FI-AAAU-2′3′-cP cleavage product. This spectrum is chimeric due to a coeluting, co-isolating molecule(s). Peaks labeled with asterisks (*) arise from coeluting, co-isolating species as determined by extracted ion chromatograms generated for the various fragment ions in the 914.14 channel. Fragment ions that have a distinct peak at 6.2 minutes were assumed to arise from the 5′-FI-AAAU-2′3′-cP cleavage product.

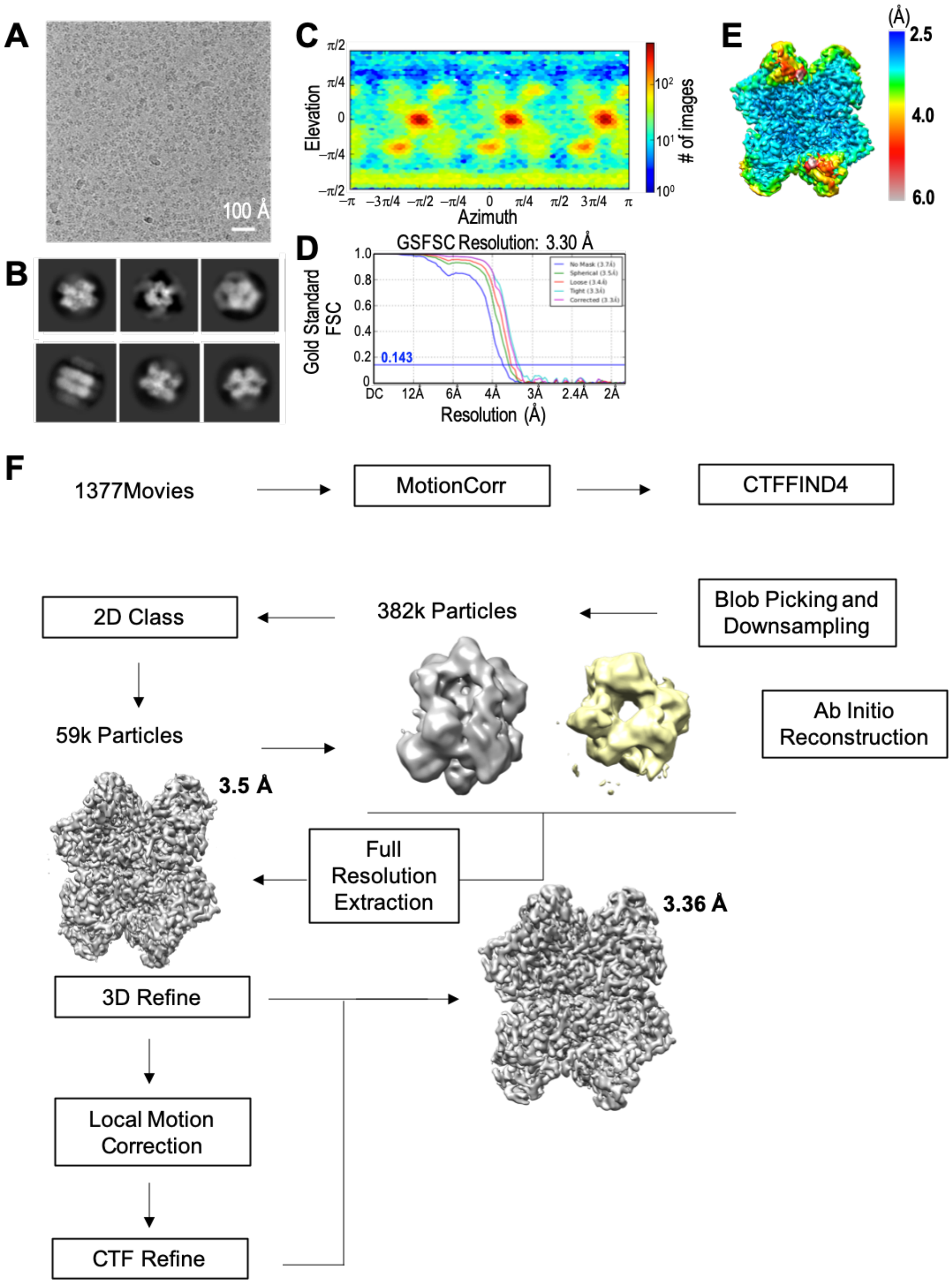

**Fig. S6. Overview of cryo-EM processing scheme for apo-state Nsp15 H235A dataset i. (A)** A representative micrograph of apo-state Nsp15 H235A in vitreous ice and **(B)** selected 2D classes generated from 1377 movies collected from an UltrAuFoil R1.2/1.3 300 mesh grid. **(C)** Angular distribution of apo-state Nsp15 H235A particles. **(D)** Fourier shell correlation (FSC) curve for the apo-state Nsp15 H235A reconstruction. The overall resolution is 3.30 Å according to the FSC 0.143 criteria (*1, 2*). **(E)** Cryo-EM reconstruction of apo-state Nsp15 H235A colored based on local resolution calculated using cryoSparc v2 (*3*). **(F)** Cryo-EM processing workflow. Picked particles (382,348) were subjected to 2D classification, 3D classification, and refinement prior to local refinement of a single Nsp15 protomer in cryoSPARC v2 (*3*).

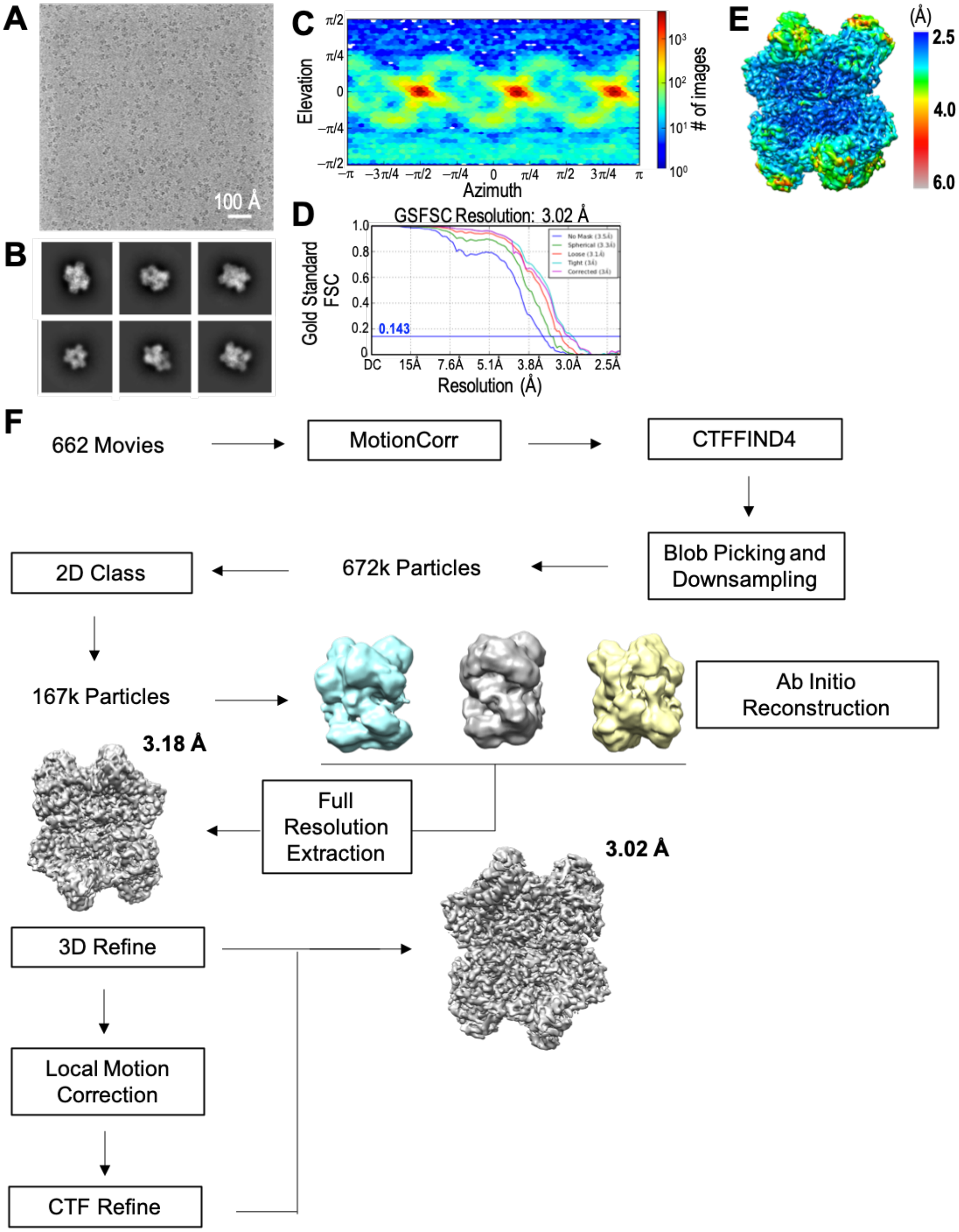

**Fig. S7. Overview of cryo-EM processing scheme for apo-state wt Nsp15. (A)** A representative micrograph of apo-state wt-Nsp15 in vitreous ice and **(B)** selected 2D classes generated from 662 movies collected from an UltrAuFoil R1.2/1.3 300 mesh grid. **(C)** Angular distribution of apo-state wt-Nsp15 particles. **(D)** Fourier shell correlation (FSC) curve for the apo-state Nsp15 reconstruction. The overall resolution is 3.02 Å according to the FSC 0.143 criteria (*1, 2*). **(E)** Cryo-EM reconstruction of apo-state Nsp15 colored based on local resolution calculated using cryoSparc v2 (*3*). **(F)** Cryo-EM processing workflow. Picked particles (672,879) were subjected to 2D classification, 3D classification, and refinement prior to local refinement of a single Nsp15 protomer in cryoSPARC v2 (*3*).

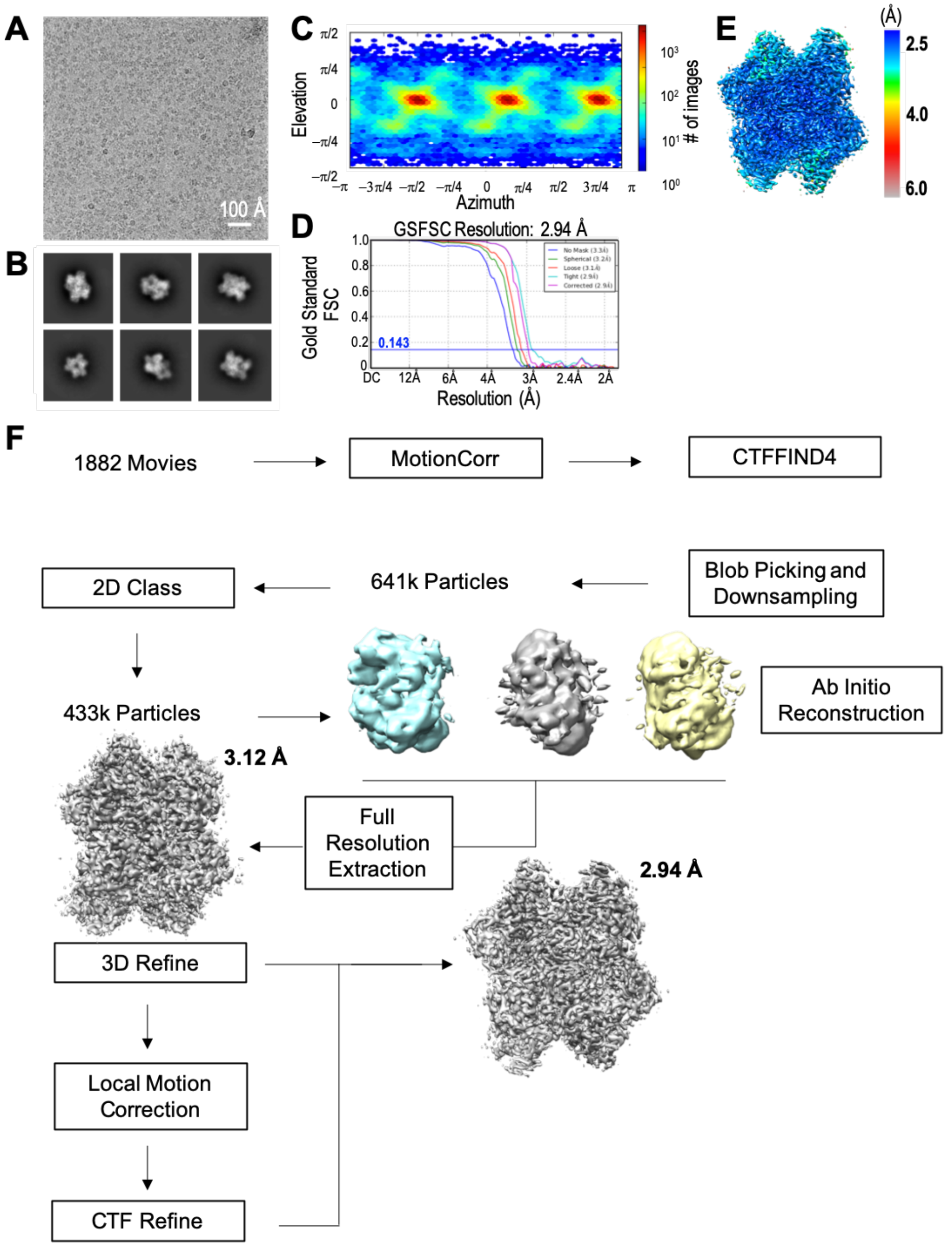

**Fig. S8. Overview of cryo-EM processing scheme for apo-state Nsp15 H235A dataset ii. (A)** A representative micrograph of apo-state Nsp15 H235A in vitreous ice and **(B)** selected 2D classes generated from 1882 movies collected from an UltrAuFoil R1.2/1.3 300 mesh grid. **(C)** Angular distribution of apo-state Nsp15 H235A particles. **(D)** Fourier shell correlation (FSC) curve for the apo-state Nsp15 reconstruction. The overall resolution is 2.94 Å according to the FSC 0.143 criteria (*1, 2*). **(E)** Cryo-EM reconstruction of apo-state Nsp15 H235A colored based on local resolution calculated using cryoSparc (*3*). **(F)** Cryo-EM processing workflow. Picked particles (641,748) were subjected to 2D classification, 3D classification, and refinement prior to local refinement of a single Nsp15 protomer in cryoSPARC v2 (*3*).

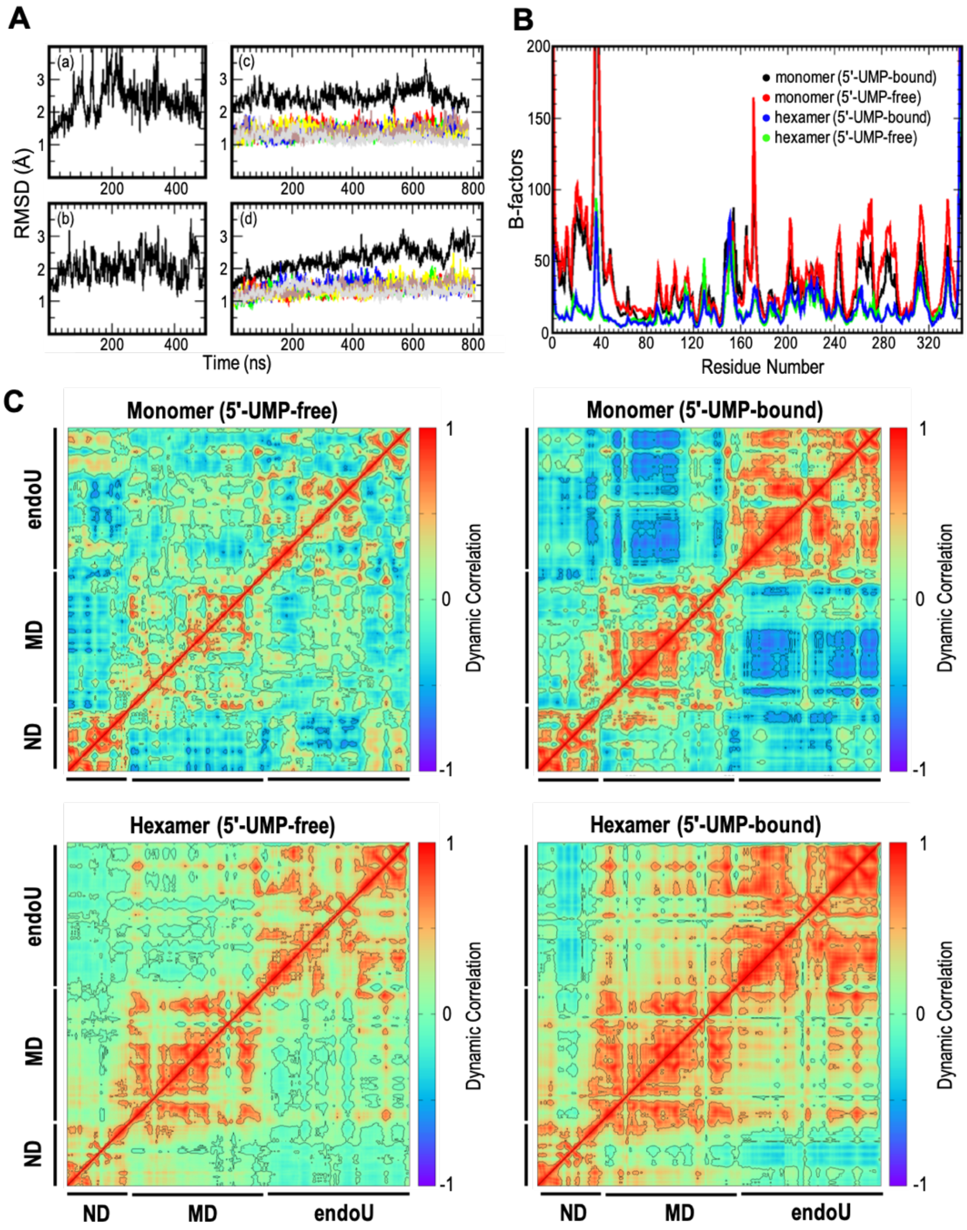

**Fig. S9. Molecular dynamics of Nsp15. (A)** Root mean square deviations (RMSDs) of (a) a substrate-free monomer, (b) 5′-UMP-bound monomer, (c) a substrate-free monomer from the hexamer (black= hexamer RMSD; colored plots= individual monomer RMSDs), and (d) a 5′-UMP-bound monomer from the hexamer (black= hexamer RMSD; colored plots= individual monomer RMSDs). Individual monomer RMSDs from hexamers systems are smaller when compared with the values of monomer systems, due to subdued dynamics of monomer in the hexameric configurations. **(B)** B-factors calculated from the positional fluctuations of the backbone atoms (red= substrate-free monomer; black= 5′-UMP-bound monomer; green= a substrate-free monomer from the hexamer; blue= a 5′-UMP-bound monomer from the hexamer). **(C)** Dynamical cross-correlation matrices (DCCM) from a 5′-UMP-free monomer, a 5′-UMP-bound monomer, a 5′-UMP-free monomer from the hexamer, and a 5′-UMP-bound monomer from the hexamer.

**Table S1**. Theoretical masses of putative Nsp15 RNA cleavage products.

| | Mass (Da) | m/z (M-H)$^-$ | m/z (M-2H)$^{2-}$ | m/z (M-3H)$^{3-}$ |
|---|---|---|---|---|
| Fl-A-3'OH | 804.215 | 803.207 | 401.100 | 267.064 |
| Fl-AA-3'OH | 1133.267 | 1132.259 | 565.626 | 376.748 |
| Fl-AAA-3'OH | 1462.319 | 1461.311 | 730.152 | 486.432 |
| Fl-AAAU-3'OH | 1768.344 | 1767.336 | 883.164 | 588.440 |
| Fl-AAAUA-3'OH | 2097.397 | 2096.389 | 1047.691 | 698.125 |
| Fl-AAAUAA-3'OH | 2426.449 | 2425.441 | 1212.217 | 807.809 |
| Fl-A-3'P | 884.181 | 883.173 | 441.083 | 293.719 |
| Fl-AA-3'P | 1213.233 | 1212.225 | 605.609 | 403.403 |
| Fl-AAA-3'P | 1542.285 | 1541.277 | 770.135 | 513.087 |
| Fl-AAAU-3'P | 1848.310 | 1847.302 | 923.147 | 615.096 |
| Fl-AAAUA-3'P | 2177.363 | 2176.355 | 1087.674 | 724.780 |
| Fl-AAAUAA-3'P | 2506.415 | 2505.407 | 1252.200 | 834.464 |
| Fl-A-2'3'-cP | 866.171 | 865.163 | 432.078 | 287.716 |
| Fl-AA-2'3'-cP | 1195.223 | 1194.215 | 596.604 | 397.400 |
| Fl-AAA-2'3'-cP | 1524.275 | 1523.267 | 761.130 | 507.084 |
| Fl-AAAU-2'3'-cP | 1830.300 | 1829.292 | 914.142 | 609.092 |
| Fl-AAAUA-2'3'-cP | 2159.353 | 2158.345 | 1078.669 | 718.777 |
| Fl-AAAUAA-2'3'-cP | 2488.405 | 2487.397 | 1243.195 | 828.461 |
| Fl-AAAUAA-TAMRA | 3294.734 | 3293.726 | 1646.359 | 1097.237 |
| 5'HO-AAAUAA-TAMRA | 2757.615 | 2756.607 | 1377.800 | 918.197 |
| 5'HO-AAUAA-TAMRA | 2428.563 | 2427.555 | 1213.274 | 808.513 |
| 5'HO-AUAA-TAMRA | 2099.510 | 2098.502 | 1048.747 | 698.829 |
| 5'HO-UAA-TAMRA | 1770.458 | 1769.450 | 884.221 | 589.145 |
| 5'HO-AA-TAMRA | 1464.433 | 1463.425 | 731.209 | 487.137 |
| 5'HO-A-TAMRA | 1135.381 | 1134.373 | 566.683 | 377.453 |
| 5'P-AAAUAA-TAMRA | 2837.581 | 2836.573 | 1417.783 | 944.853 |
| 5'P-AAUAA-TAMRA | 2508.529 | 2507.521 | 1253.257 | 835.169 |
| 5'P-AUAA-TAMRA | 2179.476 | 2178.468 | 1088.730 | 725.484 |
| 5'P-UAA-TAMRA | 1850.424 | 1849.416 | 924.204 | 615.800 |
| 5'P-AA-TAMRA | 1544.399 | 1543.391 | 771.192 | 513.792 |
| 5'P-A-TAMRA | 1215.347 | 1214.339 | 606.666 | 404.108 |

**Table S2**. Binding free energies estimated using MMGBSA calculations for the 50-ns segments of the final 450 ns of the 5′-UMP-bound Nsp15 hexamer simulation.

| Time (ns) | Nsp15 protomers of the hexameric assembly | | | | | |
|---|---|---|---|---|---|---|
| | A | B | C | D | E | F |
| 385-435 | -10.5 ± 3.7 | -23.9 ±1.9 | -18.6 ± 1.0 | -19.1 ± 1.0 | -17.7 ± 0.4 | -18.7 ± 1.0 |
| 435-485 | 4.7 ± 1.9 | -15.4 ± 0.9 | -18.3 ± 1.0 | -15.3 ± 1.4 | -19.1 ± 1.1 | -14.7 ± 1.1 |
| 485-535 | 0.0 | -5.4 ± 3.2 | -14.6 ± 1.4 | -21.3 ± 1.3 | -16.7 ± 1.1 | -18.4 ± 2.0 |
| 535-585 | 0.0 | -5.7 ± 2.8 | -1.6 ± 1,2 | -15.9 ± 1.5 | -14.8 ± 1.2 | -6.8 ± 3.2 |
| 585-635 | 0.0 | -8.7 ±3.0 | 23.2 ± 1.8 | -15.6 ± 1.4 | -13.2 ± 1.6 | -1.4 ± 2.5 |
| 635-685 | 0.0 | -7.2 ± 2.0 | 29.4 ± 1.4 | -14.2 ± 1.7 | -14.7 ± 1.4 | 4.4 ± 2.0 |
| 685-735 | 0.0 | -4.3 ± 3.2 | 34.6 ± 2.0 | -16.9 ± 1.0 | -8.4 ± 2.5 | 1.2 ± 3.1 |
| 735-783 | 0.0 | -1.2 ± 2.2 | 32.8 ± 2.6 | -17.0 ±1.3 | -6.2 ± 1.7 | -3.3 ± 2.2 |
| 785-835 | 0.0 | -0.9 ± 2.7 | 36.1 ± 2.7 | -16.7 ± 0.9 | -8.2 ± 2.8 | -1.8 ± 1.3 |

Movie S1. Cryo-EM reconstruction of nucleotide bound Nsp15.

Movie S2. 3D Variability of Apo-H235A-Nsp15 dataset i

Movie S3. 3D Variability of Apo-wt-Nsp15

Movie S4. 3D Variability of Apo-H235A-Nsp15 dataset ii
